# Flexible multitask computation in recurrent networks utilizes shared dynamical motifs

**DOI:** 10.1101/2022.08.15.503870

**Authors:** Laura Driscoll, Krishna Shenoy, David Sussillo

## Abstract

Flexible computation is a hallmark of intelligent behavior. Yet, little is known about how neural networks contextually reconfigure for different computations. Humans are able to perform a new task without extensive training, presumably through the composition of elementary processes that were previously learned. Cognitive scientists have long hypothesized the possibility of a compositional neural code, where complex neural computations are made up of constituent components; however, the neural substrate underlying this structure remains elusive in biological and artificial neural networks. Here we identified an algorithmic neural substrate for compositional computation through the study of multitasking artificial recurrent neural networks. Dynamical systems analyses of networks revealed learned computational strategies that mirrored the modular subtask structure of the task-set used for training. Dynamical motifs such as attractors, decision boundaries and rotations were reused across different task computations. For example, tasks that required memory of a continuous circular variable repurposed the same ring attractor. We show that dynamical motifs are implemented by clusters of units and are reused across different contexts, allowing for flexibility and generalization of previously learned computation. Lesioning these clusters resulted in modular effects on network performance: a lesion that destroyed one dynamical motif only minimally perturbed the structure of other dynamical motifs. Finally, modular dynamical motifs could be reconfigured for fast transfer learning. After slow initial learning of dynamical motifs, a subsequent faster stage of learning reconfigured motifs to perform novel tasks. This work contributes to a more fundamental understanding of compositional computation underlying flexible general intelligence in neural systems. We present a conceptual framework that establishes dynamical motifs as a fundamental unit of computation, intermediate between the neuron and the network. As more whole brain imaging studies record neural activity from multiple specialized systems simultaneously, the framework of dynamical motifs will guide questions about specialization and generalization across brain regions.

## Introduction

Cognitive flexibility is a key feature of the human brain. While artificial systems are capable of outperforming humans in specific complex cognitive tasks^1–3^, they so far lack flexibility for rapid learning and task switching. Therefore, a major open question in the fields of neuroscience and artificial intelligence is how the same circuit flexibly reconfigures to perform multiple tasks^4^.

Conceptual models for cognitive flexibility propose a hierarchy of elementary processes that are reused across similar tasks^5,6^. According to these models, the neural substrate for computation is modular such that combinations of previously learned subtasks may be reconfigured to perform unfamiliar tasks. This combination of subtasks is referred to as compositionality^6^. For example, a saccade task typically involves a cue that indicates in which direction to move the eyes. After learning a saccade task, a person could quickly learn an ‘anti’ version of the same task where the same cue now instructs a saccade in the opposite direction. This new task may be quickly learned by combining a computational building block for the original task with a previously learned ‘anti’ building block. These computational building blocks could be considered in task space or state space. While there is some experimental evidence that neural computation is compositional^7,8^, a concrete model for the neural implementation of compositional computation would provide major headway toward understanding cognitive flexibility for multiple tasks in the human brain.

While the time and effort required to train animals to perform many tasks has proven prohibitive to the exploration of multitask computation in biological networks, artificial neural networks now present an opportunity to explore the topic. The study of cognitive tasks through simulations in artificial networks has led to substantial advances in understanding neural computation in the past decade^9–18^. However, researchers typically trained artificial neural networks to perform single tasks in isolation, with few exceptions^19–22^, somewhat limiting the insights into biological neural circuits that perform many tasks. One exception to this trend is Yang et al. 2019^20^, in which the authors trained a single network to perform twenty related tasks and thereby identified clustered representations in state space that supported task compositionality. Yet, their analysis left open the critical questions concerning *why* and *how* recurrent networks develop a compositional organization. In this work, we identified the computational substrate that allowed for compositional computation and answered the nontrivial question of how this computational substrate is flexibly employed in different contexts.

We examined multitask networks through the lens of dynamical systems. This approach allowed us to explore the mechanisms underlying computation in a recurrently connected artificial network^23^. We found that tasks that required the same computational elements (e.g. memory, categorization, delayed-response) were implemented by sharing and repurposing dynamical motifs (e.g. attractors, decision boundaries, and rotations). We demonstrate that the dynamical motifs learned in a multitask network support compositional computation.

Dynamical motifs provided a useful granularity to study network computation. While individual network units appeared heterogenous and difficult to interpret, dynamical motifs were often interpretable and consistent across networks. Our framework provides a conceptual building block between the single artificial unit and the whole network. With the access and control afforded by the use of this simulated ‘model organism’, we explored how multiple related computations might interact in a single network. This work contributes to a better understanding of functional specialization in neural networks, and more broadly advances techniques for reverse-engineering artificial networks and analysis of high-dimensional dynamical systems.

## Results

### Network Structure

We implemented a similar input-output structure and learning protocol as in previously examined multitasking recurrent neural networks (RNNs) ^20^. These tasks included reaction-timed, delayed response and memory tasks with contextual integration, categorization, pro and anti response components (see Supp. Fig.1 and Methods Section 1.2 for task definitions). For every task, the network received three noisy inputs: fixation (1-dimensional), stimulus (4-d), and rule (15-d) (Fig.1a). The fixation input directed the network to either output zero or respond, similar to fixation cues which instruct rodents, monkeys and humans to keep the eyes and limbs still. The set of stimuli contained two separate two-dimensional vectors composed of *Asinθ* and *Acosθ*, where each vector encoded a different one-dimensional circular variable *(θ* _*1*_, *θ*_*2*_) scaled by an amplitude *A*. Depending on the rule, one stimulus vector may be contextually ignored. The rule input indicates the current task on any given trial and this information is continuously available to the network throughout each trial. Rule input was encoded in a one-hot vector where the index associated with the current task was one and all other indices were zero.

The RNN is defined by

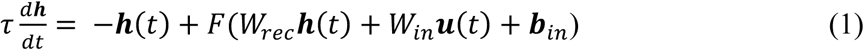

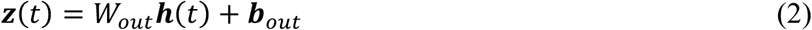

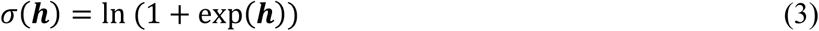

All inputs, ***u****(t) (20 × 1)*, enter the system and induce a specific pattern of activity, ***h****(t) (N*_*rec*_ *× 1)*, in the units of the RNN (Eq. 1). We refer to this *N*_*rec*_*-*dimensional vector, ***h****(t)*, as the state of the network at time *t*. The output, ***z****(t)* (3 × 1), is a linear projection of the state (Eq. 2). The output units indicate whether the network is responding in the first dimension and in which direction on a circle the RNN responds in the next two dimensions (e.g., saccade direction) (Fig.1a right). For consistency, in the majority of the paper we will focus on RNNs as described by Eq. 1, using diagonal initialization of *W*^*rec*^, the softplus nonlinear activation function (Eq. 3) and L2 activity and weight regularization. We identified shared dynamical motifs in all explored network designs, and include comparisons to other parameter choices and common network architectures throughout. All network weights were trained to minimize the squared difference between the network output and a desired (target) using back propagation through time.

Our approach to studying the trained RNNs was to uncover the underlying learned dynamical systems in order to mechanistically understand *how* the networks implement computation. This approach utilizes fixed points of Eq. 1 ^23,24^ to provide an often interpretable “skeleton” of the complex high-dimensional dynamics ^25^. By studying how fixed points change as a function of the inputs that configure the task period at hand, we may understand if and how these fixed point structures are repurposed for different computations. See Methods for further details on network setup, training, and fixed point analysis.

### Single Task Networks

To lay the groundwork to understand the multitask RNNs, we first trained individual networks to perform each task in isolation. Every task began with a context period where the rule input indicated which task to perform, the fixation signal was on, and there was no stimulus information (Fig.1a). Stimulus onset marked the beginning of the next task period (stimulus) while all other inputs remained constant. In the absence of noise, inputs to the network were piecewise constant, where every change in the inputs marked the beginning of a new task period (Fig.1a vertical lines). Therefore, during each task period with unique inputs (e.g. stimulus, context/memory, response) the network could be treated as a separate, autonomous dynamical system with a distinct set of fixed points from other task periods. Going forward, we use *dynamical motif* to mean the high-dimensional nonlinear dynamics around a fixed point skeleton that implements computation for a specified sinput. By examining locally linearized dynamics around the set of fixed points associated with a particular task period, we learn about the dynamical motifs that implement computation.

**Figure 1:**
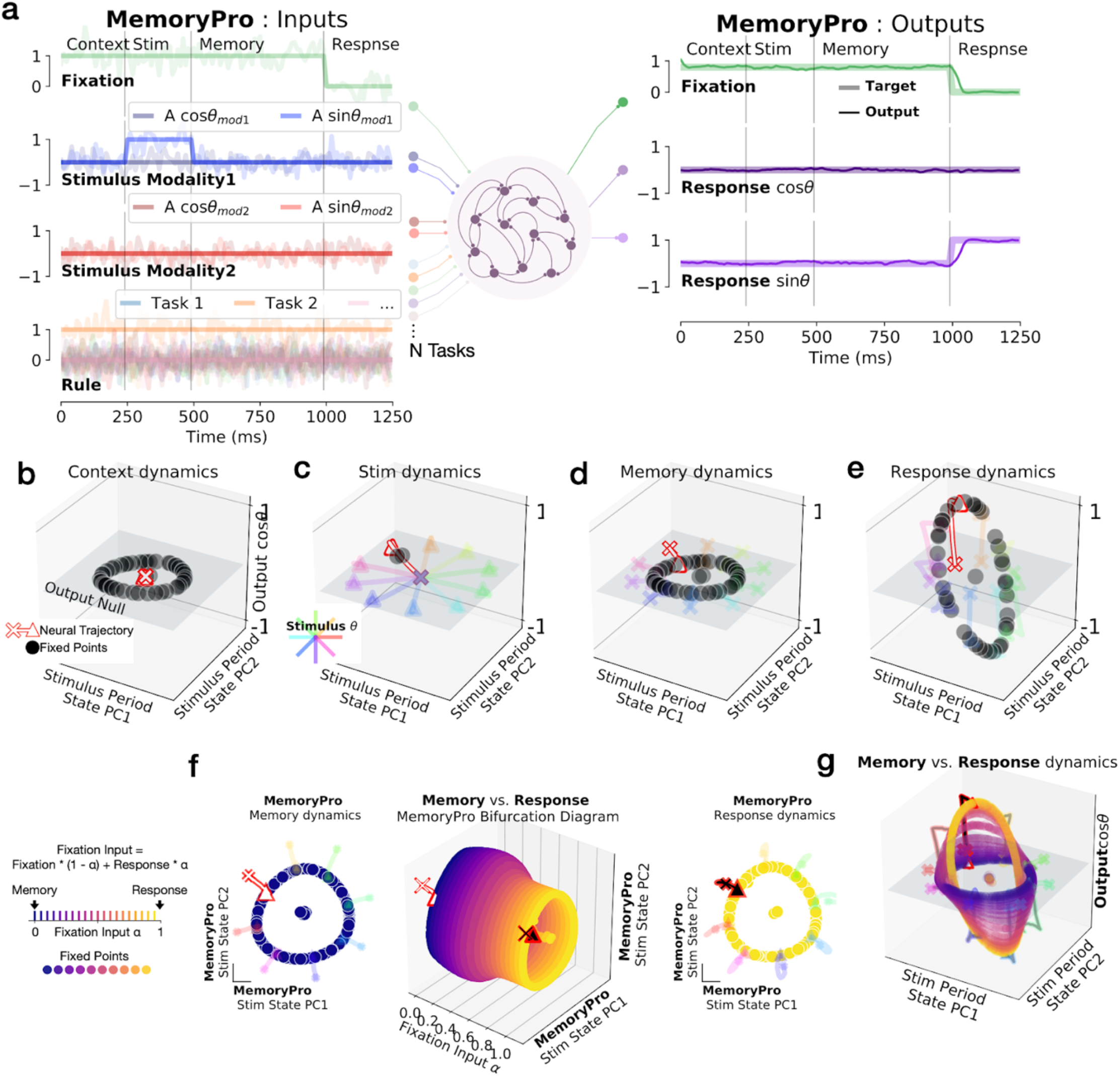
Single-task network shared fixed points across task periods. **(a) left:** Noisy fixation, stimulus (modality 1 and 2), and rule input time-series (overlayed without noise for clarity.) Noise was used during training while analyses were performed on running the network without noise. Vertical lines divide task periods: context, stimulus, memory & response. **right:** Targets (thick lines) overlaid with outputs of a trained network (thin lines). **(b-e)** State space plots for single task network performing MemoryPro task during **(b)** context **(c)** stimulus **(d)** memory **(e)** response task periods. State trajectories and fixed points projected onto the first two principal components (PCs) defined by state evolution for 100 different stimulus conditions during the stimulus period on the x and y-axes and the output weight vector (from *W*_*ou*t_) associated with *cosθ*_*stimulus*_ on the z-axis. We visualized fixed point locations for *θ*_*stimulus*_*=0* (black dots) in all subpanels of Figure 1 and additionally plotted state trajectories for other stimulus conditions (see Methods Section 1.5 for further details on fixed point identification). State trajectories (colored lines) colored according to stimulus orientation with *θ*_*stimulus*_*=0* highlighted in red, starting from ‘x’ and ending with ‘▴’. **(f-g)** Interpolation between inputs for memory (*α=0*) and response (*α=1*) periods **(f) middle:** Fixed points for 20 intermediate *α* values (x-axis) projected into top two stimulus period PCs (as in b-e) (y and z-axes) with (**left)** memory *α=0* and **(right)** response *α=1* fixed points and trajectories. **(g)** Fixed points for input-interpolation between memory (blue) and response inputs (yellow). State trajectories colored according to stimulus orientation. Same axes as (b-e).

For example, in a MemoryPro task (respond in the same direction as the stimulus after a memory period), there were 4 periods (visually divided by vertical lines Fig.1a). In the first period (context), the rule input indicated which task the network performed for that trial. In a network trained to perform only the MemoryPro task, the context period inputs result in one fixed point at the center of a ring of fixed points (Fig.1b). The central fixed point serves as an initial condition for performing the task computation during the ensuing stimulus period. Notice the context period inputs are identical to the memory period (rule and fixation on, stimulus off) (Fig.1a), so the fixed points are necessarily identical between these task periods. We will show later that the additional ring of fixed points was relevant to the memory computation during the memory task period.

In the stimulus period, we examined the fixed point structure for each stimulus input separately. Stimulus period state trajectories for different stimuli diverged from the central initial condition towards stimulus-dependent fixed points, mapping out a stimulus representation that was orthogonal (null) to the response readout dimension (Fig.1c). While fixed points were identical in the context and memory periods, the network state interacted with different fixed points due to different initial conditions. During the memory period, the state evolved toward a ring of fixed points that made up an approximate ring attractor (locally attracting structure in all dimensions except tangent to the ring, which is neither contracting nor expanding) (Fig.1d). Together, these fixed points stored the identity of the stimulus orientation based on the initial conditions of the state at the beginning of the memory period (end of the stimulus period). Finally, in the response period, the fixation input changed to zero and a new ring attractor emerged (Fig.1e). During the response period, the ring was oriented such that it had a non-zero dot product with the output weights (*W*_*out*_) and was therefore output potent^26^. The new ring caused the network to respond in the appropriate orientation based on the initial conditions of the response period (end of the memory period). Thus, the network responded with the appropriate orientation for this task (*φ*_*response*_*=θ*_*stimulus*_).

What is the relationship between the ring of fixed points in the memory and response periods? To address this question, we traced locations of the fixed points as we interpolated across memory and response period inputs, *(1-α) u*_*memory*_ *+ α u*_*response*_ with *α* in 0.05 increments between 0 and 1. We identified fixed points for each incremental input setting as a function of *α* (see Methods Section 1.6). We employ this method frequently and call it input-interpolation fixed point identification (shortened to input interpolation). By interpolating across input conditions for the memory and response periods, we traced how fixed points moved and possibly changed stability as the dynamical system reconfigured for different task periods.

We interpolated the fixation input from its memory period value (1) to its response period value (0). For every intermediate input value throughout this interpolation, an approximate ring attractor was present (Fig.1f). The smooth transition of this fixed point structure implies that each intermediate ring attractor was functionally the same ring attractor across input conditions. In this single task network, the dynamical motif that performed memory and response computations was shared across task periods. The ring attractor rotated from output null space into output potent space when the fixation input changed to zero (Fig.1g). The change in inputs across task periods shifted the location of the fixed points and thus changed their collective computational purpose.

### Two Task Networks

We then simultaneously trained networks to perform two tasks on interleaved trials. We studied two-task networks to learn if and how the addition of a second task changed the fixed point structures compared to single task networks. The MemoryPro and MemoryAnti tasks were both memory guided response tasks that received identical stimulus inputs. The target outputs in the pro task were the same as the stimulus inputs (*φ*_*response*_*=θ*_*stimulus*_) whereas in the anti task, targets were in the opposite direction as the stimulus (*φ*_*response*_*=θ*_*stimulus*_*+π*) (see Supp. Fig.1 and Methods Section 1.2 for full task definitions).

Input-interpolation across rule inputs for a network trained on the MemoryPro and MemoryAnti tasks revealed shared fixed points across tasks during the context/memory, stimulus and response periods (Fig.2). Context period fixed points were similar to the single task network throughout rule input interpolation, with one stable fixed point that was relevant to the context period and a ring of fixed points that was relevant to the memory period (Fig.2a-b,e-f). Stimulus period rule input interpolation revealed two separate stable fixed points and an unstable fixed point between the basins of attraction for each intermediate input condition (Fig.2c-d). The network state evolved away from the unstable fixed point, which smoothly moved in state space across interpolated input conditions, resulting in the network state evolving toward a different stable fixed point for each task. From that point onward, the state interacted with a shared ring attractor across both tasks (MemoryPro and MemoryAnti) and task periods (memory and response) according to the response direction (Fig.2e-h). This network flexibly performed two related computations through small changes in fixed point locations. In addition to shared fixed points across different tasks and task periods, we could identify shared fixed points across different stimulus conditions for the same task period (Supp Fig.2a-c). In this work, we will explore how these changes in the location of shared fixed points enable a single network to perform different computations and why this structure results in a compositional organization of tasks in multitask networks.

**Figure 2.**
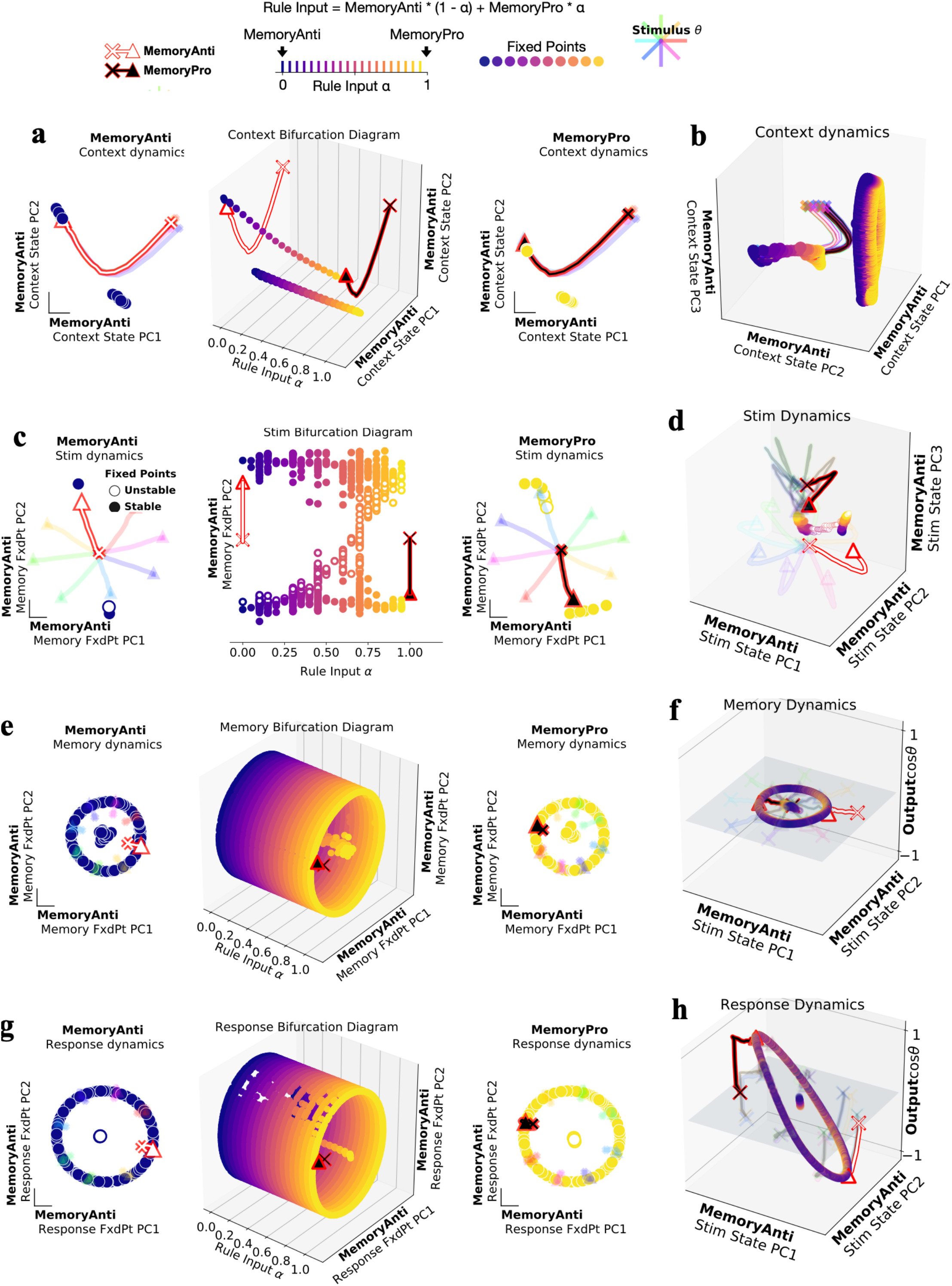
Two-task networks shared fixed points across related tasks. Fixed points for interpolation between inputs for MemoryAnti (*α=*0) and MemoryPro (*α=*1) tasks during **(a**,**b)** context **(c**,**d)** stimulus **(e**,**f)** memory and **(g**,**h)** response periods. **(a) middle:** Fixed points for 20 intermediate *α* values (x-axis) projected into top two PCs defined by state evolution during the context period of the MemoryAnti task (y and z-axes) with (**left)** MemoryAnti *α=0* and **(right)** MemoryPro *α=1* fixed points and trajectories. **(b)** Fixed points for rule input-interpolation between tasks, MemoryAnti (blue fixed points, white state trajectory) and MemoryPro (yellow fixed points, black state trajectory) projected into the top three MemoryAnti context period state evolution PCs. **(c)** Same as a for stimulus period, with unstable (open) and stable (closed) fixed points projected into top PCs defined by fixed points during the memory period of the MemoryAnti task (y-axis) **(d)** Same as b for stimulus period, projected into the top three MemoryAnti stimulus period state evolution PCs. **(e)** Same as a for memory period, projected into top two PCs defined by fixed points during the memory period of the MemoryAnti task (y and z-axes) **(f)** Same as b for memory period, projected into the top two MemoryAnti stimulus period state evolution PCs (x and y-axes) and the output weight vector (from *W*_*out*_) associated with *cosθ*_*stimulus*_ on the z-axis. **(g)** Same as a for response period, projected into top two PCs defined by fixed points during the response period of the MemoryAnti task (y and z-axes) **(h)** Same as f for response period.

One might expect that networks share fixed points due to the limited computational resources in small networks. We therefore trained networks that were nearly an order of magnitude larger and without noise to determine how abundant computational resources might change this solution. To our surprise, we found even large networks without noise still shared dynamical motifs (see Supp Fig.2d-k). We interpret these findings to mean shared dynamical motifs are not a product of limited resources and rather provide a more universal solution for multitask computation.

### Identifying Dynamical Motifs in 15 Task Networks

To quantify shared structure across a large number of tasks in a single network and also to compare this shared structure across multiple networks, we developed a modified version of the task variance metric described by Yang et al. 2019 ^20^. We were motivated to study task periods because changes in the inputs reconfigure the RNN’s dynamics across different task periods. For example, when the stimulus input turns off in some tasks, the network goes from processing a stimulus to maintaining a memory of the stimulus. Aside from training noise, the inputs are static within a task period and therefore the RNN’s dynamics are fixed. Task periods, therefore, provide the relevant granularity to identify the dynamical motifs that perform distinct computations.

We divided tasks into task periods and computed the variance across stimulus conditions for each unit, normalized across all task periods (see Methods Section 1.9). The result of this variance analysis was a matrix of each unit’s normalized variance for each task period of every task (Fig.3a), which we refer to as the variance matrix in subsequent analyses. We sorted the rows and columns of this matrix based on similarity to better visualize its inherent structure (see Methods Section 1.10).

**Figure 3.**
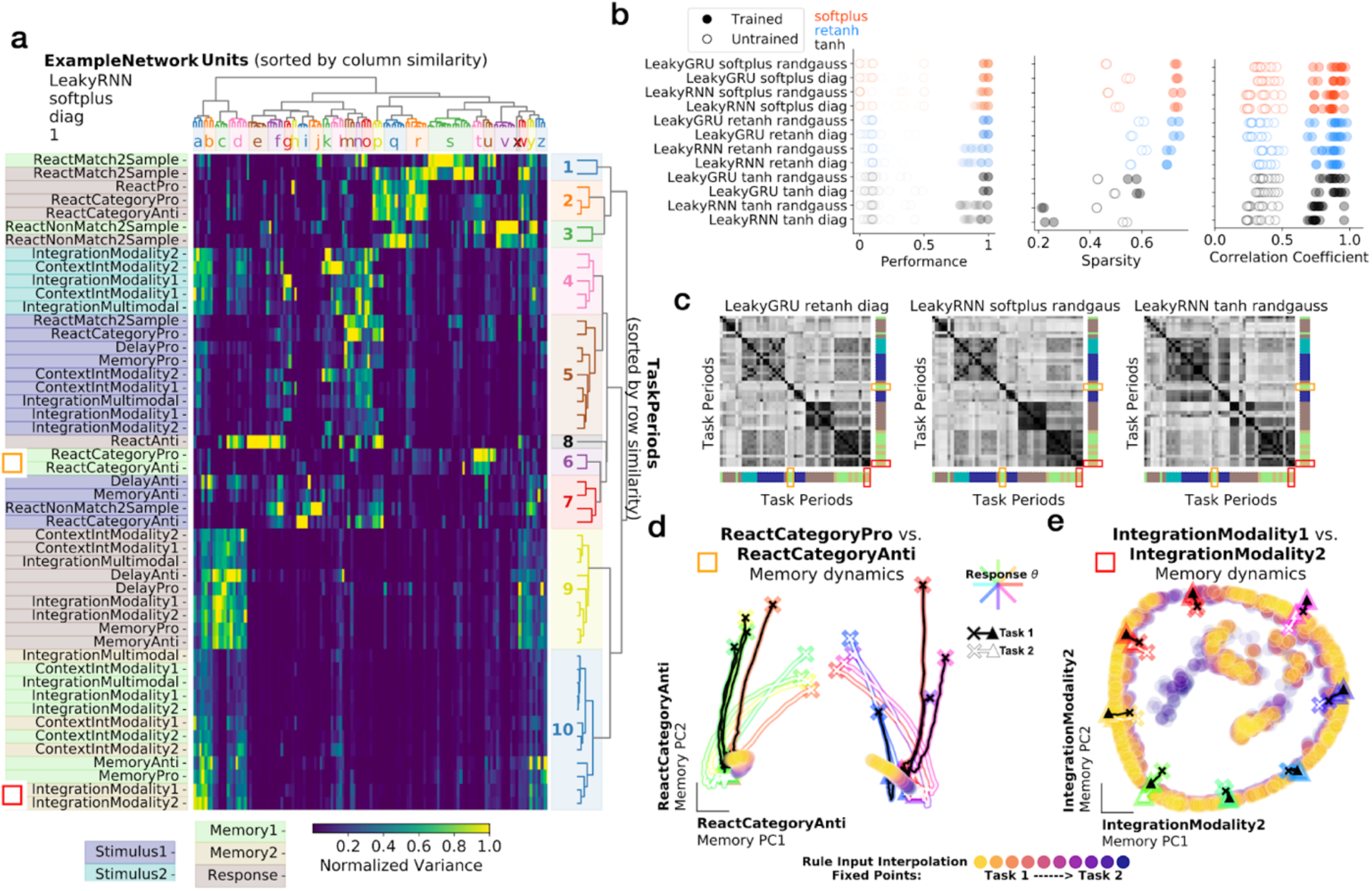
Modular organization in 15 task networks was not dependent on activation function or network initialization. **(a)** Variance matrix: variance of unit activations across stimulus conditions, normalized across task periods (columns normalized by the maximum entry in each column). Rows and columns sorted according to similarity (see Methods Section 1.10). **(b) left:** Performance across all tasks for 3 networks of each of 12 hyperparameter settings. **middle:** Sparsity of the task variance measured as the fraction of entries >15% maximum unit variance. **right:** Task period correlation matrix (examples shown in **c**) for trained and untrained networks are sorted according to rows in a and correlated to trained networks for all other hyperparameter settings. **(c)** Correlation matrix of rows in variance matrix (as in a) for 3 different example networks; rows and columns sorted according to rows in a. Orange and red rectangles highlight discrete memory (ReactCategoryPro, ReactCategoryAnti) and continuous memory (IntegrationModality1, IntegrationModality2) tasks respectively. **(d)** Shared point attractors for two category memory tasks as seen by input interpolation across tasks during memory period. State trajectories for 8 stimulus conditions (colored by response direction) starting from ‘x’ in ReactCategoryAnti state evolution PC space for ReactCategoryPro (black) and ReactCategoryAnti (white) tasks. Rule input interpolation across tasks during memory period with fixed points for intermediate rule input conditions in filled circles. **(e)** Same as d for two continuous circular variable memory tasks, highlighting shared ring attractors. State trajectories starting from ‘x’ in IntegrationModality2 state evolution PC space for IntegrationModality1 (black) and IntegrationModality2 (white) tasks.

The variance matrix can be considered as a map of functional specialization in the network, similar to the BOLD signal in an fMRI scan, or widefield imaging data. This analysis could be performed directly on neural data to learn about the computational substructure for multiple tasks in a behaving animal. Sorting the rows and columns of the variance matrix revealed a blockwise structure, where groups of units had large variance for groups of task periods with similar computational requirements (Fig.3a). Similar computations can be seen in the task period color labels (*Stimulus1, Stimulus2* - for tasks with two sequential stimulus presentations, *Memory, Response*, etc) and in the task names (Category, DecisionMaking, Memory, etc) (Fig.3a, left). For example, task period cluster #2 (Fig.3a, right) corresponds to reaction timed response task periods (see Supp. Fig.1 for definitions of all tasks). These tasks receive new stimulus information during a response period that must be incorporated into the computation immediately. Therefore, the network cannot prepare a response direction before the fixation cue disappears. On the other hand, in task period cluster #9, the network receives no new information during the response period and must instead use the memory of the stimulus to produce the correct output during the response period. These separate blocks in the variance matrix reveal two distinct clusters of units that contribute to response period dynamics: one for tasks with reaction timed responses and another for tasks with memory guided responses. Other unit clusters for stimulus (unit cluster k-o, task period cluster #4-5) and memory (unit cluster a-b, task period cluster #10) computations are apparent in the block-like structure aligned with task period type (Fig.3a task titles and task period color labels to the left of the variance matrix).

Block structure in the variance matrix was robust to different network architectures and hyperparameters (examples in Supp. Fig.3). To quantify task period similarity across networks with different design choices, we calculated the variance matrix for each trained network. We then sorted task periods according to similarity of rows in one reference network and computed the correlation matrix of the sorted rows in the variance matrix for each network (see Methods Section 1.9). The correlation matrix for each trained network revealed the block wise similarity of task periods (Fig.3c). By comparing the correlation matrices across networks (Pearson correlation coefficient between correlation matrices, see Methods Section 1.9), we found that trained networks were more correlated to every other trained network than to untrained networks (Fig.3b right). Higher correlation across trained networks compared to untrained networks confirmed the block structure in the variance matrix emerged from learning the task computations rather than from network design choices or the structure of the inputs.

As we will show, the clusters of the variance matrix arise from shared dynamical motifs across task periods. This shared structure takes the form of attractors, decision boundaries and rotations. We highlight two different examples of shared memory dynamical motifs using rule input interpolation (Fig.3d-e) and highlight their positions in the variance matrix (Fig.3a left of task period names, red and yellow squares). A pair of memory category task periods are within the same cluster in the variance matrix, suggesting their computations are performed by a similar set of units (Fig.3a yellow square left: task period cluster #6, unit cluster t). Both category tasks utilized the same two point attractors for memory of the initial stimulus. Rather than store the identity of the initial continuous circular stimulus, the network stored which category it must respond to, regardless of task (Fig.3d). In another example of a shared attractors across tasks, we found a ring attractor that was shared across several tasks (Fig.3e, Fig.3a task period cluster #9-10, unit clusters a-d). All of these tasks required the memory of the initial continuous circular stimulus variable. To show that this ring attractor was shared across tasks, we interpolated across rule inputs for a pair of these tasks (IntegrationModality1 and IntegrationModality2) and found a similar shared ring structure as in the two task networks (Fig.2e). We highlight shared category and continuous memory dynamical motifs in networks with different activation functions in Supp. Fig.4.

In addition to clusters of task periods with similar variance, there are also some task periods that do not cluster with other task periods. These can be seen in the variance matrix as isolated row segments with high variance and in the correlation matrices as rows with low correlation across all other task periods. For example, task period cluster #8 is dedicated to the ReactAnti task, cluster #1 is dedicated to the ReactMatch2Sample task and cluster #3 is ReactNonMatch2Sample (Fig.3a). In these cases, the computation performed in the unique task period is so distinct from other computations the network performs, the dynamical motif is not reused across tasks. The set of tasks that employed unique dynamical motifs was similar across hyperparameter settings (Fig.3c, Supp. Fig.3).

### Motif Alignment to Unit Axes

One notable difference across network hyperparameters was sparsity in the variance matrix. We define sparsity to be the fraction of entries in the variance matrix below a threshold of 15% maximum unit variance. Networks with non-negative nonlinear activation functions had sparse task variance matrices whereas networks with the *tanh* activation function, which has a range of (−1, 1) did not (Fig.3b middle). We understand this sparsity to be a function of optimal network performance requiring potentially interfering dynamical motifs to be organized into orthogonal subspaces. In a network where all units can take only positive values, this orthogonalization can only occur along unit axes (Supp. Fig.5), thus resulting in sparse variance matrices for networks with non-negative activation functions. We find clusters to be present in *tanh* networks, simply not aligned to unit axes and therefore non identifiable using methods described in Yang et al. 2019^20^. By examining the correlation matrix and the correlation coefficient across networks, we see that similar clusters are present in the *tanh* networks (Fig.3b right, c right).

### Shared Stimulus Period Dynamical Motifs in 15 Task Networks

The variance matrix provides a useful overview of which task periods are implemented by similar clusters of units, but falls short of addressing exactly *how* these subpopulations implement shared dynamical motifs. Shared motifs are implemented by organizing the state in the appropriate region of state space to evolve on the relevant shared dynamical landscape. To walk through this explanation in detail, we focus on stimulus period dynamics and highlight two examples, one in which dynamical motifs are shared and another where motifs are not shared.

Tasks with similar stimulus computations (noisy integration, pro vs. anti, reaction-timed vs. delayed response) organized their initial conditions for the stimulus period to be nearby in state space and evolved in a similar way after stimulus onset (see schematic Fig.4a). We visualized this organization of stimulus period initial conditions in principal component (PC) space defined by the final state of the context period across all tasks (Fig.4b). To summarize the relationship between initial conditions and the ensuing stimulus dynamics for different tasks, we compared pairs of trials presented with the same stimulus across different tasks. We plotted the Euclidean distance between initial conditions against the angle between the state evolution on the first time step for these pairs of trials (Fig.4c). We observed that pairs of tasks with similar computations had initial conditions that were closer together and had smaller angles between state trajectories on the first time step of the stimulus period compared to pairs of tasks with distinct computations. Similar initial conditions for stimulus onset resulted in shared context dependent stimulus amplification in some networks (Supp Fig.6). In these cases, the state update was scaled in magnitude according to whether the stimulus input was either modality one or two, dependent on the position of that state at stimulus onset.

**Figure 4.**
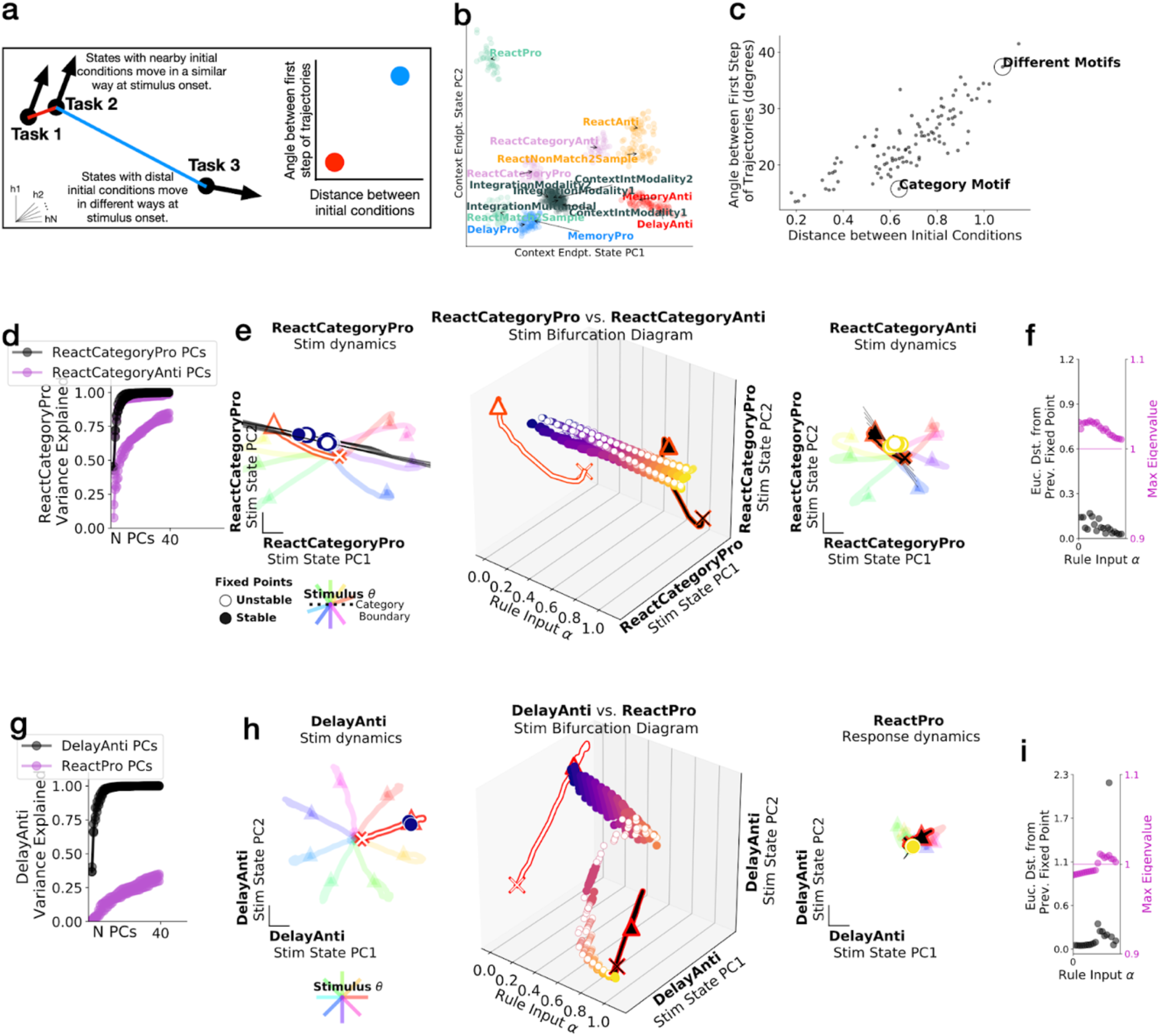
Tasks with similar stimulus computations were in nearby parts of state space and shared dynamical motifs. **(a)** Schematic of analyses in b and c. **(b)** The state for each trial (colored dot for each of 20 trials on each task) at the end of the context period (just before stimulus period) projected onto the top two PCs defined by the state at the end of the context period for all tasks. Trials colored by similar stimulus computations as given by task definitions: delayed response pro (blue), delayed response anti (red), noisy integration (gray), categorization (pink) reaction-timed pro (teal) and reaction-timed anti (orange). **(c)** Euclidean distance between pairs of trials from different tasks at the end of the context period plotted against cosine angle between same pair after stimulus onset for a particular stimulus input for one timestep, then averaged across stimulus angle inputs. Pairs of tasks in d-i circled and labeled, **(d-f) ‘Category Motif’:** ReactCategoryPro and ReactCategoryAnti, **(g-i) ‘Different Motifs’:** DelayAnti and ReactPro **(d)** Fraction of variance explained for ReactCategoryPro task by the ReactCategoryAnti task PCs (purple) compared to its own PCs (black) for 5 trained networks with different random seeds. **(e)** Rule input interpolation across category tasks for one stimulus angle. **middle:** Unstable (open) and stable (closed) fixed points for 20 intermediate *α* values (x-axis) projected onto top two PCs defined by state evolution during the stimulus period of the ReactCategoryPro task (y and z-axes) with (**left)** ReactCategoryPro *α=0* and **(right)** ReactCategoryAnti *α=1* fixed points and trajectories for 8 different stimulus angles (rainbow colors) Expanding dimensions around unstable fixed points are visualized as black lines. **(f)** Euclidean distance between fixed points (black) and maximum real eigenvalue for the linearization of the state update around each fixed point (purple) for the single unstable fixed point closest to the state at the end of the stimulus period for 20 consecutive *α* values between 0 and 1. We analyzed only one unstable fixed point that is most proximal to the end of the state trajectory for each input condition (see Methods Section 1.7). **(g-f) ‘Different Motifs’:** DelayAnti and ReactPro. Same as (d-i), but for DelayAnti and ReactPro tasks. **(i)** Euclidean distance between fixed points (black) and maximum real eigenvalue for the linearization of the state update around each fixed point (purple) for the single fixed point closest to the state at the end of the stimulus period for 20 consecutive *α* values between 0 and 1. We analyzed one fixed point that is most proximal to the end of the state trajectory for each input condition.

The relationship between context period states and stimulus period trajectory angles across tasks support the idea that nearby initial conditions allowed tasks with similar stimulus computations to reuse the same dynamical landscape and therefore evolve in similar ways. To provide further detail, in (Fig.4d-i) we examine these features in two examples of comparisons between tasks that (1) share the same stimulus period dynamical motif and then tasks that (2) do not share the same dynamical motif.

In the case of the two categorization tasks, ReactCategoryPro and ReactCategoryAnti, we found a shared stimulus motif (Figure 4d-f). In the ReactCategoryPro task, the network was trained to respond if both sequential stimuli were less than or both greater than π; whereas, in the ReactCategoryAnti task the network was trained to respond if stimuli were on opposite sides of π. In either task, there was a decision boundary at *θ*_*stimulus*_ *= π*. The initial conditions for these tasks were nearby and trajectories during the stimulus period were aligned (Fig.4c ‘Category Motif’). We quantified stimulus response overlap by computing the fraction of variance explained for the evolution of the state trajectory on one task by the other task’s PCs (purple) compared to its own PCs (black) (Fig.4d), revealing that both tasks were performed in an aligned subspace. Aligned stimulus responses for both category tasks were visualized in PC space defined by the stimulus period state trajectories of the ReactCategoryPro task (Fig.4e left, right). These analyses revealed that activity evolved in a qualitatively similar way for trials with the same stimulus conditions on both tasks, suggesting the state evolution could have occurred on a similar dynamical landscape.

To better understand the relationship between the dynamical landscapes across task contexts, we interpolated across rule inputs during the stimulus period for both category tasks with the same stimulus input. We visualized the fixed points for each intermediate rule input configuration (Fig.4e middle). We found that similar stimulus responses were governed by shared stable and unstable fixed points, demonstrated by the smooth bridge of fixed points between both tasks. We projected the unstable dimension of each unstable fixed point into PC space and found this dimension was aligned with the direction of the state evolution and roughly orthogonal to the decision boundary. (Fig.4e). We defined the most relevant fixed point to be the closest unstable fixed point to the state at the end of this task period. This simplification of one relevant fixed point was often necessary to tease apart how relevant dynamics are reconfigured across tasks while several to hundreds of other fixed points related to computations during other task periods also moved through state space. A continuous bridge mapped movement of the relevant fixed point, suggesting that rule inputs shifted a relevant shared fixed point that was reused across both tasks (Fig.4e). We quantified the distance between consecutive locations of the relevant fixed point for each interpolated input, highlighting the smooth and small transition of the fixed point location across tasks (Fig.4f). Moreover, the stability of the local linear dynamics around this shared fixed point was consistent across all intermediate input conditions, as shown by the the maximal real part of the eigenvalue of the linearized RNN state update around each interpolated fixed point location (see Methods Section 1.7) (Fig.4f). This real part of the eigenvalue does not pass below the threshold of 1, where the fixed point would become stable. We interpret this result to mean that both category tasks reuse the unstable fixed point that moves the state away from the category boundary.

The DelayAnti and ReactPro tasks were an example pair that did not share any dynamical motifs (Fig.4g-i). The DelayAnti task began with the context period, followed by a stimulus presentation that signaled the opposite response direction (*φ*_*response*_*=θ*_*stimulus*_ *+π*), followed by a go cue that signaled when to initiate a delayed response (see Supp. Fig.1 for all task definitions). The ReactPro task began with the context period, followed by a stimulus presentation that signaled the same response direction and required an immediate response. During the context period, the network state evolved toward dissimilar locations for trials of either task and trajectories during the stimulus period were not aligned (Fig.4c ‘Different Motifs’). We defined the subspace for the ReactPro task by performing principal components analysis on the state trajectories during the stimulus period. We projected the DelayAnti task in the same subspace and found that very little variance was captured by the other task PCs (Fig.4g). We also visualized both tasks in a subspace defined by the first two PCs of the DelayAnti task and again found very little overlap, suggesting both tasks evolved in mostly non-overlapping subspaces (Fig.4h right). We interpolated across these two rule inputs during the stimulus period, revealing that there was a bifurcation where the relevant stimulus-dependent fixed point did not form a continuous bridge across interpolated rule inputs (Fig.4h middle). We quantified the distance between consecutive fixed points that were closest to the endpoint of the state trajectory for each interpolated input and identified a large discrete jump in the location of the relevant fixed point (Fig.4i). We visualized the maximal real part of the eigenvalues of the linearized RNN state update around each consecutive fixed point, revealing dissimilar local dynamics around fixed points for interpolated input conditions (Fig.4i).

Taken together, these features suggest that shared dynamical motifs are implemented by evolving the state to the appropriate region of state space such that it interacts with a shared fixed point across similar task computations. Category tasks shared both unstable and stable fixed points. On the other hand, stimulus period dynamics for the DelayAnti and ReactPro tasks evolved in separate subspaces and were governed by different stable fixed points. These analyses revealed that shared structure was not merely an artifact of all tasks within the same network. Rather only tasks with similar computations implemented shared dynamical motifs. All subspace and distance analyses in this section could be experimentally tested on neural data of animals performing multiple tasks.

### Dynamical Motifs Result in Modular Lesion Effects

The authors of Yang et al. 2019^20^ previously found that network lesions affected sets of tasks that shared computational features. For example, if the output of a particular cluster of units was set to zero, then all tasks involving a particular computation decreased their performance, while other tasks were unaffected. Their work left open the major question of *why* lesion effects were modular. We identified the cause of these modular lesion effects to be related to the underlying modular dynamical motifs that perform computation.

We examined the impact of lesioning clusters of units described in the variance matrix in Fig.3a. We lesioned a cluster of units by setting the output of all units within the cluster to be zero throughout a given trial. Clusters were identified by performing hierarchical clustering on the columns and rows of the variance matrix and identifying a distance criterion to maximize the ratio of intercluster to intracluster distances, which resulted in clusters of units a-z (Fig.3a, Methods Section 1.10).

Many unit clusters had high variance for a set of task periods with similar computations. For example, unit clusters a and c had high variance for memory and response task period clusters 10 and 9 respectively (Fig.3a). Other unit clusters had high variance for modality 2 stimulus periods (unit cluster k), or anti stimulus periods (unit cluster f), etc.

In two example lesions, we demonstrate that each unit cluster lesion impacted only a subset of tasks that shared computational features where units had high variance, reproducing a finding previously reported in Yang et al. 2019. Here, we show that lesions either did or did not impact task-relevant computations depending on whether the relevant underlying dynamical motif was impacted (Fig.5). A lesion to one cluster only impacted performance on tasks that included a delay period (Fig.5a,b,e). We visualized state trajectories for a task that did include a delay period (MemoryPro) and a task that did not include a delay period (ReactPro) in lesioned (red) and non-lesioned networks (blue)(Fig.5b). State trajectories were dramatically impacted when the delay-related dynamical motif was lesioned during the performance of the MemoryPro task, but not for the ReactPro task. In a second example, we lesioned a cluster of units with high variance during the performance of anti-response tasks and found there was little to no change in the performance or state trajectories of pro-response tasks (Fig.5c,d,f).

**Figure 5.**
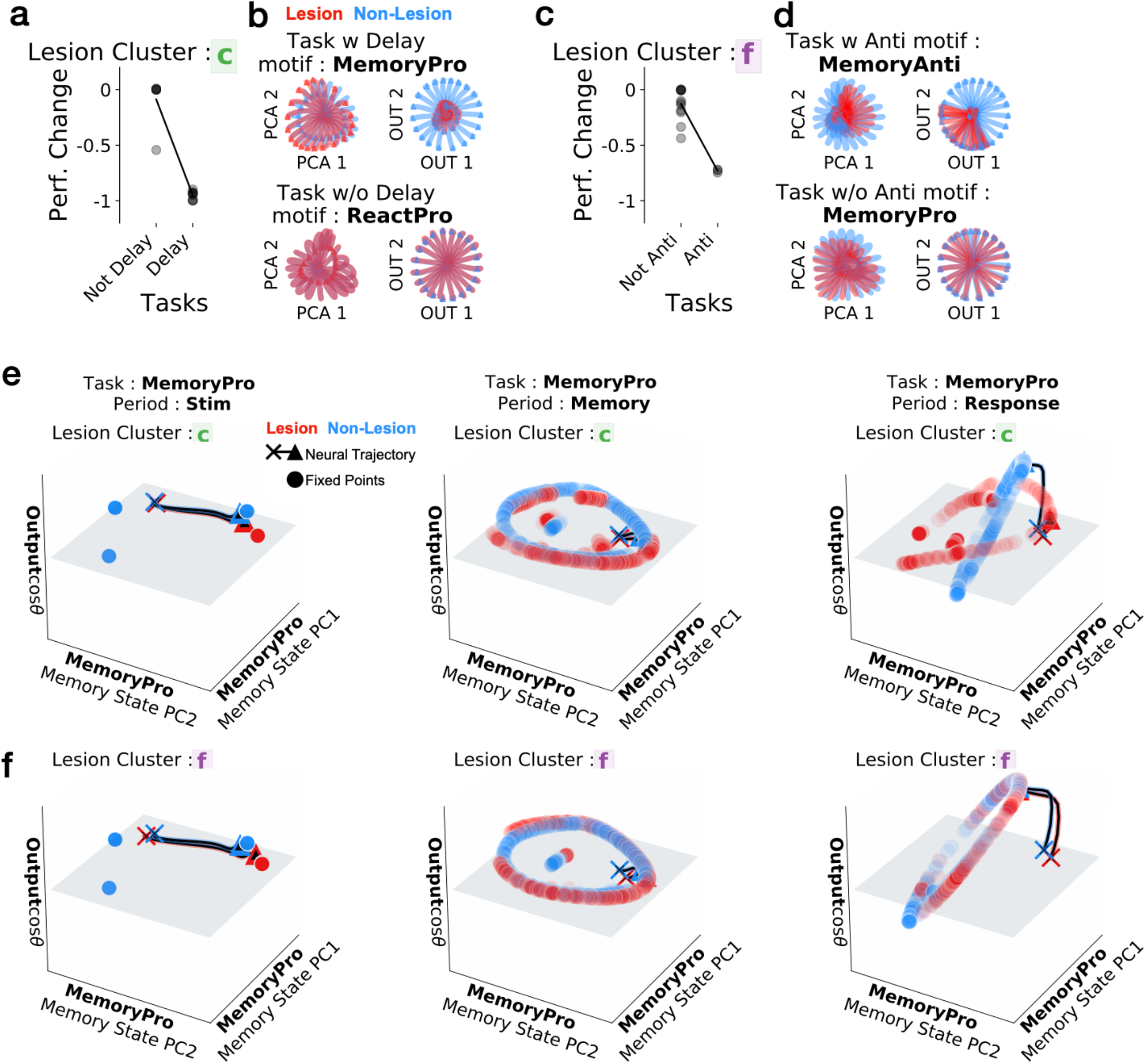
Unit cluster lesions had modular effects on task period clusters that shared the same dynamical motif. **(a**,**c)** Fraction performance change after lesion (by setting unit output to zero) to a cluster of units with high variance on **(a)** delayed-response (cluster c in Fig.3a) **(c)** anti-response (cluster f in Fig.3a) relevant task periods. **(b**,**d)** State evolution in **(left)** PC space defined by the state evolution for the non-lesioned network and **(right)** output weight space (from *W*_*out*_) for **(top)** task with relevant computational feature (i.e. delay period, anti-response) and **(bottom)** task without relevant computational feature for lesioned (red) and non-lesioned (blue) trials. **(e)** Fixed points and state trajectories for *θ*_*stimulus*_*=0* with a lesion to cluster c (red) and non-lesioned (blue) networks (same as in panel a,b) during **(left)** stimulus, **(middle)** memory and **(right)** response periods for MemoryPro task projected onto the first two PCs (x and y-axes) defined by state evolution during the memory period of the MemoryPro task without lesion and the output weight vector (from W_out_) associated with *cosθ*_*stimulus*_ on the z-axis. State trajectories in black emanating from ‘x’ and ending with ‘▲’. **(f)** Same as e with lesion to cluster f.

When a unit cluster was lesioned, all task periods that shared the relevant dynamical motif were impacted. When tasks did not utilize the dynamical motif impacted by a unit cluster lesion, they were not impacted by the lesion. For example, MemoryPro stimulus period activity was not impacted by a lesion to the unit cluster which implemented the memory and response period dynamical motif of the same task (Fig.5e left). The state evolved toward a stable fixed point in approximately the same location in both the lesioned (red) and full network (blue) after this cluster was lesioned. However, this lesion resulted in discontinuity of the relevant ring attractor in the memory and response periods and minimal rotation into output potent space during the response period (Fig.5e middle, right). Conversely, there was little change in the fixed point structure after a lesion to the anti-stimulus motif (cluster f) for all task periods during the MemoryPro task (Fig.5f). Taken together, results from our lesion studies suggest that modular lesion effects are a result of modular fixed point structures that implement dynamical motifs.

### Fast Learning of Novel Tasks by Reusing Dynamical Motifs

Networks were able to rapidly learn new tasks sequentially by reconfiguring previously learned dynamical motifs. We first identified a task where each task period shared a dynamical motif with at least one of the other 14 tasks: MemoryAnti. In this task, the network uses the anti stimulus motif (as shown in two task network, Fig.2c) and the delayed-response memory motif (as in one and two task networks Fig.1f, 2e,g). We next trained a network to perform every task except the MemoryAnti task. After learning all input, recurrent and output weights and biases for other tasks, we trained only the one-hot rule input weights to learn the MemoryAnti task. This vector of length *N*_*re*c_ maps a single rule input onto the recurrent weights (Fig.6a). By only training the single rule input weights, we did not interfere with any previously learned dynamical motifs that were constructed in the recurrent weight matrix, *W*^*rec*^, enabling learning of new tasks without catastrophic forgetting. The network was able to learn the MemoryAnti task when previously trained on all other tasks (Fig.6b, black).

To determine if the anti stimulus motif and delayed response motif are sufficient for this effect, we pre-trained a new network on a key set of tasks that included these motifs: DelayAnti and MemoryPro. These networks could also sequentially learn the MemoryAnti task with single rule input training (Fig.6b, blue). Pre-training on this minimal set of tasks with relevant dynamical motifs learned as fast and to similar proficiency as networks pretrained on all tasks, suggesting the relevant dynamical motifs were sufficient for this effect. Conversely, if we pretrained the network on two pro tasks that did not include the anti motif (orange) or if we started from a network with no pre-training (green), networks required many more training steps or could not learn the MemoryAnti task at all (Fig.6b).

Without pre-training on any tasks that included the anti computation, the network had to learn a new dynamical motif by modifying only the *N*_*re*c_-dimensional rule input vector. This resulted in stimulus period state trajectories that were not highly overlapping with previously learned tasks (Fig.6c). The MemoryAnti stimulus period relevant fixed point was distinct from the previously learned DelayPro stimulus period relevant fixed point (Fig.6e,f). However, the network was pre-trained on a task that included a relevant memory period dynamical motif, the MemoryPro task. Despite learning a new stimulus period anti motif, the network was still able to reuse the previously learned memory motif. The state converged on the previously learned ring attractor (Fig.6g-j). This result highlights the modularity of dynamical motifs.

Networks that were pre-trained with the relevant dynamical motifs reused the anti-stimulus and memory dynamical motifs for fast learning of the novel MemoryAnti task. We found that MemoryAnti state trajectories were in highly overlapping subspaces with the DelayAnti task during the stimulus period (Fig.6k-l) and with the MemoryPro task during the memory period (Fig.6o,p). Rule input interpolation between both anti tasks during the stimulus period (Fig.6m) and memory tasks during the response period (Fig.6q) provided strong evidence that the fixed point structures were shared.

Tasks with unique motifs that were not shared with other tasks could not be learned as well using this pre-training method compared to full network training. We trained networks on all tasks except one, for each task, and then trained the rule input weights for the held out task. We compared the training cost of these networks to single task networks where all weights were trained on only the held out task (Supp Fig.7a,b).

We wanted to better understand the relationship between how well the pre-training method works for a given task and the uniqueness of its relevant dynamical motifs. We identified tasks with unique response period motifs using our task period variance matrix (Fig.3a), as rows with low correlations to other rows (task periods with low correlation to other task periods) (See Methods Section 1.12 for details). Three response task periods that were dissimilar from all other response periods could not be learned as well by rule input training alone

(Supp Fig.7c). In summary, we found that rapid learning was not possible in the context of novel tasks with unique dynamical motifs. These results provide support that rapid learning of novel tasks required reconfiguration of relevant previously learned dynamical motifs.

## Discussion

In this work, we addressed the question of *how* recurrently connected artificial networks flexibly repurpose their learned dynamics to perform multiple tasks. Our collection of commonly studied cognitive tasks could be broken down into an underlying set of subtasks (contextual integration, memory, categorization, anti-response, etc). We showed that networks learned this underlying subtask structure, which resulted in specialized computational building blocks that we call dynamical motifs, dedicated to each subtask. Using input interpolation and fixed point analyses, our work showed how dynamical motifs were organized in relation to one another and often shared across tasks or task periods. Inputs reconfigured the dynamical system in each task period, resulting in often smooth changes to the dynamical landscape underlying the performed computation. The motifs necessary to perform each subtask included different types of attractor structures, input amplifications, decision boundaries and rotations. This modular subtask structure in our set of tasks is analogous to the structure of language, algebraic thought and other natural behaviors in everyday life ^27,28^.

Our framework of examining subtask computation through the lens of dynamical motifs made it possible to explain lesions and learning results described previously ^18,20^. As in Yang et al. 2019, we found that lesioning specific unit clusters resulted in specific deficits in sets of tasks that were related computationally. Units within a cluster had high variance during a set of task periods that shared a dynamical motif. When we lesioned a given unit cluster, the fixed points that made up the associated dynamical motif were greatly impacted in terms of their locations and stability. A unit cluster associated with one dynamical motif could be lesioned with little impact to other computations the network performed. That is, disturbances in the fixed point structure of one motif had little impact on the fixed point structures that implemented other dynamical motifs. This finding was surprising given the all-to-all connectivity possible in our networks, as well as the fact that no regularizations or constraints to induce modularity were employed in the training of the recurrent neural networks. Recent work on subpopulation structure for the implementation of multiple complex tasks provides insight for these findings ^21,29^.

We demonstrated that networks equipped with relevant dynamical motifs were able to repurpose those motifs in a modular fashion for fast learning of novel tasks. The initial phase of learning novel dynamical motifs was a slow process. However, given a rich repertoire of previously learned motifs, a network could quickly repurpose motifs to perform novel tasks by learning a single input vector. We found that training with modifiable recurrent weights resulted in faster and higher performance learning of novel dynamical motifs than training input weights alone (Fig.6b, orange, green). However, sequential task learning with changes to the recurrent weights leads to catastrophic forgetting ^18,30^. Our findings suggest a useful lifelong learning strategy could include two stages of learning. Early in learning, it may be beneficial for highly plastic recurrent connections throughout the brain to learn novel subtasks (dynamical motifs). Late in learning, reduced plasticity in recurrent connections and new plastic layers that function as contextualizing inputs could repurpose previously learned subtasks. This hypothesis is interesting to consider in the context of critical periods ^31^ and re-aiming ^32^. We hypothesize that this two-stage process of slow and fast learning could provide some intuition for off-and on-manifold brain-machine interface learning results in nonhuman primates ^33–35^. Additionally, the ability to repurpose dynamical motifs may inform our thinking about state-of-the-art models that require pre-training ^36^.

Our results are based on examining artificial systems, which are missing many of the constraints and complexities of biological neural circuits. We did not consider diverse cell types or prescribed architectures in our networks, and only applied noise-corrupted static inputs. We expect our results to inform a larger class of computational systems, while future work is required to link our findings directly to biological systems. More broadly, artificial networks have proven to be a valuable new tool for generating hypotheses about possible solutions to computation in biological networks ^9–18^. While our learning rules are not biological, we hypothesize that optimized artificial neural networks and the principles we uncover from them are informative about biological neural circuits based on principles of optimality and robustness ^37^. While some constraints changed the *way* dynamical motifs were shared, our main finding that they *are* shared was robust across all types of networks and hyperparameter choices that we tested, including large networks without noise (Supp Fig.2d-k). These findings suggest shared motifs are not a result of limited computational resources. We hypothesize that the modular, compositional organization of dynamical motifs was a result of the modular subtask structure of our tasks but learning dynamics through gradient descent could play a role. It will be of great interest to further explore the prevalence of dynamical motifs in other artificial^38^ and biological systems^5,39^.

Dynamical motifs underlying modular computation were organized into largely distinct subspaces (Fig.3a, 4g, 6c). This organization of dynamical motifs allowed for both modular lesion effects and rapid learning through the composition of distinct motifs without interference. In networks with nonlinearities that constrained activations to be positive, dynamical motifs were aligned to unit axes, highlighting these largely distinct subspaces.

Fixed point structures often moved in different contexts rather than appear, disappear or change stability (Fig.1f, 2, 3d-e). For example, we found that during the stimulus task period, a stable fixed point moved in state space according to the stimulus input orientation (Supp Fig.2a), resulting in the same qualitative fixed point structure for each stimulus orientation. We found that multitasking networks sometimes employed parameter bifurcations across input conditions for dissimilar computations. The relationship between high dimensional parameter bifurcations, composition and computation is an important area of future research; see for example recent work ^40–43^.

Fixed points often persisted as we varied the inputs, even when the associated motif was not relevant to the computational task at hand (Fig.2c). These irrelevant fixed points did not interfere with network computation because the network state was organized to be more proximal to the task relevant fixed points. This is related to the idea of sloppiness in under-constrained systems ^44–46^.

Shared dynamical motifs can be thought of as both modular and continuous. For our broad set of hyperparameters, dynamical motifs were modular in our networks due to the modularity of subtasks within our defined collection of tasks (contextual integration, memory, categorization, anti-response, etc). If our set of tasks was made up of a continuous spectrum of related tasks (e.g. contextual integration of two input modalities weighted by a continuous parameter), we might expect to find a dynamical motif that smoothly varies as the continuous parameter is varied (e.g. a left eigenvector of a line attractor that rotates in accordance with the weighting). Smoothness of dynamics was a prevalent feature in our networks, where similar computations were implemented in nearby regions of state space, on a similar dynamical landscape (Fig.4a-c). This feature of smoothness is consistent with biological noise-robust networks ^13,16,47^.

Our findings resulted in a number of experimentally testable results. The method of studying unit variance across tasks ^20^, which we expanded to task periods to identify units contributing to dynamical motifs, could be readily performed on neural data (Fig.3a). This analysis could be informative about perturbations to biological network activity that would most dramatically impact performance on a computation of interest (Fig.5). Our findings could apply to biological networks on multiple spatial scales. For example, we could consider multiple motifs implemented within a cortical region or we could think of different cortical areas as implementing different motifs. Our approach for training networks sequentially by reusing previously learned dynamical motifs could be used to determine ideal curricula for training animals on complex tasks. For example, given a particular task of interest, one could train an artificial network to perform the task and inspect all relevant dynamical motifs. For example, the ‘anti’ and ‘memory’ motifs were the sufficient set of relevant motifs in Figure 6b. Based on the task relevant motifs, one could systematically design a set of tasks to learn a sufficient set of motifs rather than designing a curriculum through guesswork. Additionally, this work highlights the relevance of reporting training protocols as they likely shape the dynamical motifs that implement computation^48,49^. Beyond experimental predictions, our work provides some intuition for why we find functional specialization in the brain.

**Figure 6.**
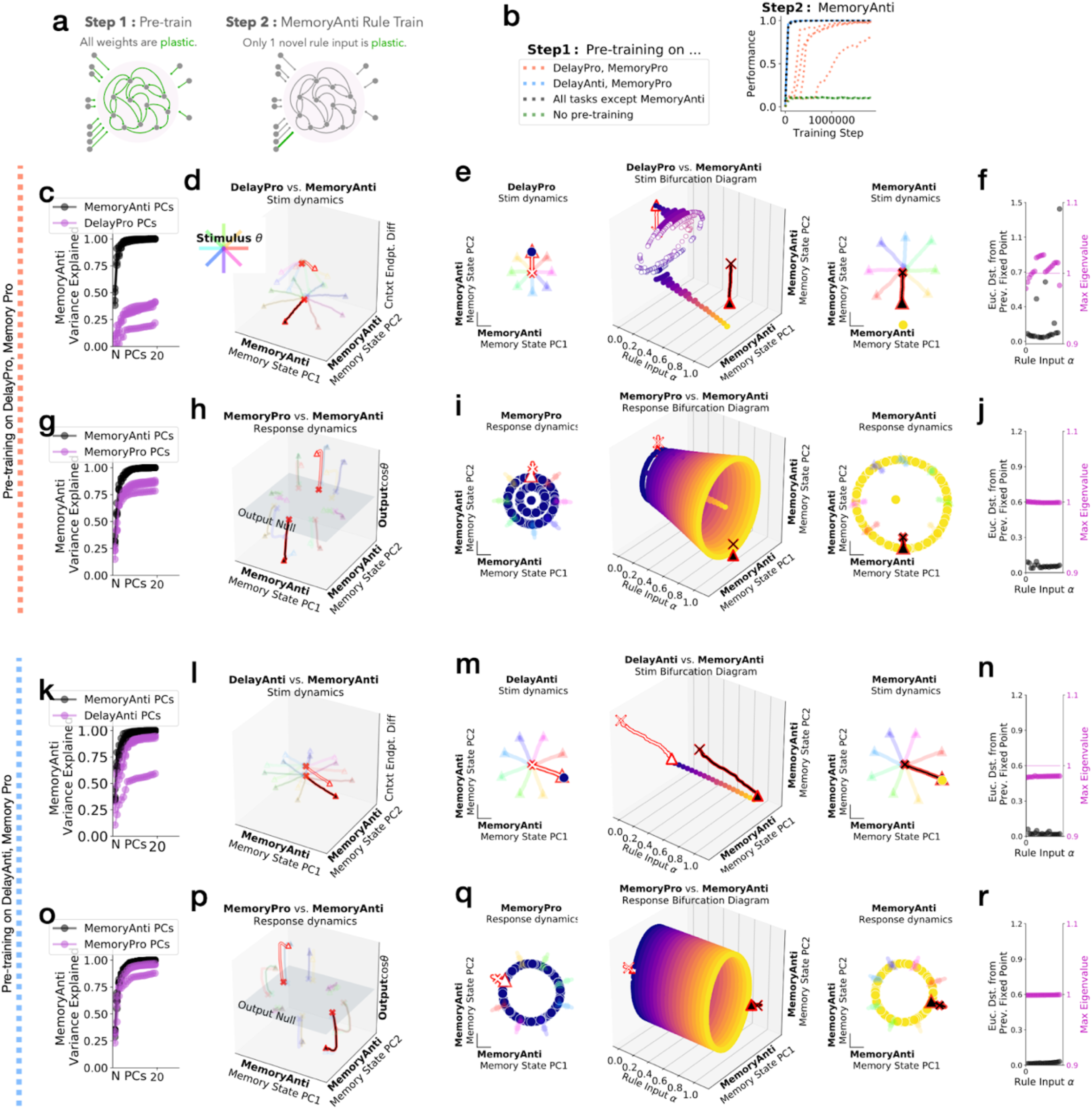
Dynamical motifs were reused for fast learning of novel tasks with familiar computational elements. **(a)** Schematic of two stage learning. Networks were pre-trained on a set of tasks while all weights were plastic. The same network was then trained on a novel task by only learning a single one-hot rule input. **(b) left:** Networks were pre-trained on two tasks that include pro and memory motifs (orange), anti and memory motifs (blue), all motifs (black), no motifs (green). **right:** Performance during MemoryAnti task rule input training after pre-training on various sets of motifs for five different networks each. **(c-j)** Network pre-trained on DelayPro and MemoryPro tasks (pro and memory motifs) was then trained to perform MemoryAnti through weight changes to the MemoryAnti rule input (length *N*_*rec*_ vector). **(c)** Fraction of MemoryAnti stimulus period variance explained by MemoryAnti stimulus period PCs (black) and DelayPro stimulus period PCs (purple) quantifies the extent to which both pro and anti tasks are in a similar subspace during the stimulus period. **(d)** Stimulus period state trajectories for DelayPro (white) and MemoryAnti (black) tasks for 8 different stimulus angles (rainbow colors) projected into PC space (x and y axes) defined by state trajectories for 100 different stimulus angle inputs during MemoryAnti task and context period state endpoint difference between both tasks (z-axis). **(e)** Rule input interpolation across tasks for one stimulus angle. **middle:** Unstable (open) and stable (closed) fixed points for 20 intermediate *α* values (x-axis) projected onto top two PCs defined by state evolution during the memory period of the MemoryAnti task (y and z-axes) with (**left)** DelayPro *α=0* and **(right)** MemoryAnti *α=1* fixed points and trajectories for 8 different stimulus angles (rainbow colors) **(f)** Euclidean distance between fixed points (black) and maximum real eigenvalue for the linearization of the state update around each fixed point (purple) for the single fixed point closest to the state at the end of the stimulus period for 20 consecutive *α* values between 0 and 1. Only analyzing one fixed point that is most proximal to the end of the state trajectory for each input condition (see Methods Section 1.7). **(g-j)** same as c-f for response period of MemoryPro and MemoryAnti tasks. **(k-r)** Network pre-trained on DelayAnti and MemoryPro tasks (anti and memory motifs) was then trained to perform MemoryAnti task through weight changes to the MemoryAnti rule input (length N_rec_ vector). **(k-n)** same as c-f for stimulus period of DelayAnti and MemoryAnti tasks with pre-training on DelayAnti and MemoryPro tasks. **(o-r)** same as c-f for response period of MemoryPro and MemoryAnti tasks with pre-training on DelayAnti and MemoryPro tasks.

In summary, through the lens of dynamical systems, we identified the underlying computational substrate for clustered representations described previously in Yang et al. 2019^20^ and highlighted a new level of organization between the unit and the network: groups of units that implement dynamical motifs. More broadly, our findings highlight the relevance of dynamical systems as a framework to better understand the response properties of neurons in the brain. As researchers record more whole brain activity, the framework of dynamical motifs will guide questions about specialization and generalization across brain regions.

## Data Availability and Code Availability

All trained networks and analysis code will be published online before the time of publication.

## Acknowledgements

We thank Lea Duncker, Niru Maheswaranathan, Dan O’Shea, Matt Golub, Frank Willett and Larry Abbott for feedback on this manuscript. We also thank our larger scientific community for feedback on this work over the years. The authors acknowledge support by Simons Collaboration on the Global Brain (SCGB) project numbers 543045 and 543049.

## Author Contributions

LD, KS and DS conceived of this project. LD performed analyses under the supervision of KS and DS. LD, KS and DS wrote the manuscript.

## Competing Interests

KS serves on the Scientific Advisory Boards (SABs) of MIND-X Inc. (acquired by Blackrock Neurotech, Spring 2022), Inscopix Inc. and Heal Inc. He also serves as a consultant / advisor (and was on founding SAB) for CTRL-Labs (acquired by Facebook Reality Labs in Fall 2019, and is now a part of Meta Platform’s Reality Labs) and serves as a consultant / advisor (and is a co-founder, 2016) for Neuralink. DS works for Meta Platform’s Reality Labs, but the work presented here was done entirely at Stanford. LD has no competing interests.

Correspondence and requests for materials should be addressed to.

## 1 Methods

### 1.1 Network Structure

We examined “vanilla” continuous-time RNNs for the majority of this work, although see methods subsection on varying hyperparameters and architectures [Section 1.4]. Before time discretization,

RNN network activity **h**, a vector of length *N*_*rec*_, followed the dynamical equation

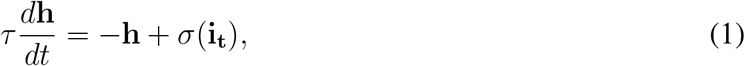

with the total neuron input **i**_**t**_ defined as

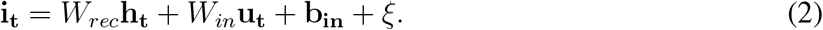

*W*_*in*_ and *W*_*rec*_ were the input and recurrent connection matrices of size *N*_*rec*_ × *N*_*in*_ and *N*_*rec*_ × *N*_*rec*_. These matrices specified the contribution of the inputs and upstream network activity to downstream network activity. The bias vector, **b**_**in**_, was of length *N*_*rec*_. The private noise variable, *ξ*, was *N*_*rec*_ independent Gaussian white noise processes with zero mean and standard deviation of 0.05.

The state of this system evolved over time according to the current state **h** and inputs to the system **u**. The nonlinear function *σ*(.) was chosen to be softplus, tanh or retanh [see Section 1.4 for details]. The time constant, *τ*, specified the rate of decay of the network state. After using the first-order Euler approximation with a time-discretization step ▵*t*, we had

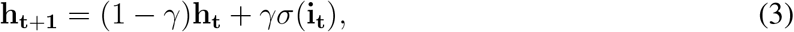

where *γ* ≡ Δ*t*/*Τ*, which we set to 0.2. The full update equation was

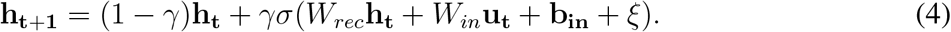

A set of output units **z** were read out from the network according to

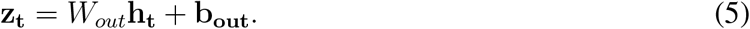

*W*_*out*_ was the output connection matrix of size *N*_*out*_ × *N*_*rec*_ and **b**_**out**_ was a bias vector of length *N*_*out*_. All *W* matrices, *W*_*in*_, *W*_*rec*_, *W*_*out*_ and bias vectors **b**_**in**_, **b**_**out**_ were learned over the course of training [see Section 1.3 for details].

The network received four types of noisy input.

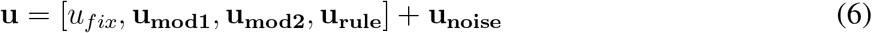

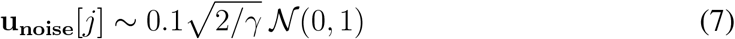

The fixation input *u*_*fix*_ was 1 when the network was required to fixate and 0 when the network was required to respond. The stimulus inputs **u**_**mod1**_ and **u**_**mod2**_ each a length-2 vector of (*A* sin *θ* and *A* cos *θ*) representing a different “modality” and each modality representing a one-dimensional circular variable described by the degree around a circle. The strength of the stimulus inputs varied in amplitude according to *a*. We greatly reduced the dimensionality of the stimulus inputs from the original implementation of these tasks in order to simplify visualizations and analysis. This simplification in stimulus inputs required removal of five of the original 20 tasks, because we could no longer present multiple stimuli in the same modality simultaneously. The network also received a set of rule inputs encoded in the vector, **u**_**rule**_. This vector represented which task the network was supposed to perform on each trial as a one-hot vector. The rule input unit corresponding to the current task was 1, while other rule input units were 0. Therefore, the number of rule input units was equal to the number of tasks trained. The rule unit activation patterns for different rules were orthogonal to each other in this 1-hot encoding. Therefore, relationships between tasks were learned by the network rather than baked into the inputs. Finally, each input had Gaussian noise added to it according to equations (6-7). Here the input noise strength was scaled by the factor 0.1. Note this was an order of magnitude greater than in previous work, to prevent over-fitting [1].

In total, there were

*N*_*in*_ = 1(fixation) + 2(modalities)× 2(*A* sin *θ* and *A* cos *θ*) + 15(rule) = 20 input units.

The network projected the state, **h**_**t**_, to an output ring, which contained 2 units (sin *ϕ*, cos *ϕ*) to encode response direction on a circle. In addition, the network projected **h** to a fixation output unit, which should be at the high activity value of 1 before the response and at 0 once a response is generated.

In total, there were

*N*_*out*_ = 1(fixation) + 1(modality)× 2(sin *ϕ* and cos *ϕ*) = 3 output units.

### 1.2 Tasks and performances

Inputs and outputs for an example trial on each task are shown in Supplementary Figure 1. Fixation input was 1 for the duration of the trial until the response period, when fixation input changed to zero. Reaction timed tasks never received a go cue; therefore, the fixation input was always at 1 and the network was required to break fixation to respond as soon as the relevant stimulus arrived. Target fixation output activity was high 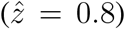 before the response period and low 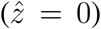 during the response period for all tasks. If the activity of the fixation output prematurely fell below 0.5, the network was considered to have erroneously broken fixation and the trial was incorrect. The response direction of the network was read out in a 2 dimensional vector (sin *ϕ* and cos *ϕ*). The decoded response direction was considered correct if it was within π/10 of the target direction.

Tasks could be divided into periods, where each task period was a segment of sequential time steps with continuous inputs (without noise). Each set of inputs reconfigured the network into a new dynamical landscape, with a different fixed point structure. Distinct dynamical landscapes for each input condition is a crucial concept for this work and should be emphasized. For all tasks, in the first period (context) the rule input provided the network with information about task context. The onset of stimulus information marked a change in the stimulus inputs and the beginning of the next task period (stimulus1). All tasks had at least one stimulus period, but some had two stimulus periods. The period between the stimulus and response or between two stimuli was the memory period (memory1). If there was a second stimulus (stimulus2), sometimes the network was required to respond immediately to the second stimulus and in other tasks there was an additional memory period (memory2) before the response period (response). The duration of the context, stimulus1, memory1, stimulus2, memory2, and response periods were *Tcontext, Tstimulus*1, *Tmemory*1, *Tstimulus*2, *Tmemory*2, *Tresponse*, respectively. We adjusted the distribution of task periods to be wider than in previous work and drawn from a uniform distribution to prevent the network from predicting task period transitions. These modifications had a simplifying effect on fixed point structures).

*U* (*t*_1_, *t*_2_) is a uniform distribution between *t*_1_ and *t*_2_. The unit for time is milliseconds. Stimuli were presented in either modality 1 or 2 at random unless stated otherwise.

#### Delayed Response

(2 tasks) DelayedPro: Move in same direction as stimulus (*ϕ*_*response*_ = *θ*_*stimulus*_) after delay. DelayedAnti: Move in opposite direction as stimulus (*ϕ*_*response*_ = *θ*_*stimulus*_ +*π*) after delay. Stimulus remains on throughout stimulus and response periods.

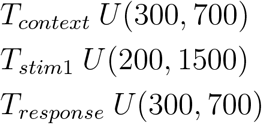

#### Memory Response

(2 tasks) MemoryPro: Move in same direction as stimulus (*ϕ*_*response*_ = *θ*_*stimulus*_) after memory. MemoryAnti: Move in opposite direction as stimulus (*ϕ*_*response*_ = *θ*_*stimulus*_ + *π*) after memory. Stimulus disappears for memory period and remains off during response.

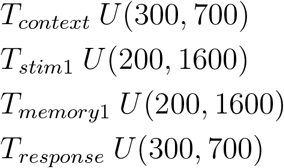

#### Reaction Timed

(2 tasks) ReactPro: Move in same direction as stimulus (*ϕ*_*response*_ = *θ*_*stimulus*_) immediately. ReactAnti: Move in opposite direction as stimulus (*ϕ*_*response*_ = *θ*_*stimulus*_ + *π*) immedi-ately.

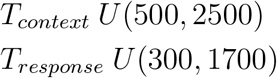

#### Decision Making

(5 tasks) Move in direction of stimulus with largest amplitude. IntegrationModal-ity1: Only modality 1 is presented. IntegrationModality2: Only modality 2 is presented. ContextInt-Modality1: Both modalities presented, only attend modality 1. ContextIntModality2: Both modalities presented, only attend modality 2. IntegrationMultimodal: Both modalities presented, attend both modalities equally.

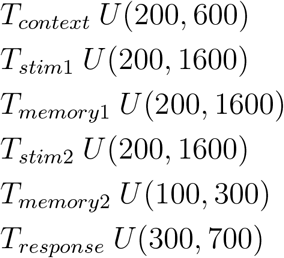

#### Delay Match

(4 tasks) Immediately move in direction of *θ*_*stim*2_ if sequentially presented pair match. ReactMatch2Sample: Match if same angle (*θ*_*stim*1_ = *θ*_*stim*2_). ReactNonMatch2Sample: Match if opposite angle (*θ*_*stim*1_ = *θ*_*stim*2_ + *π*). ReactCategoryPro: Match if same category (*θ*_*stim*1_, *θ*_*stim*2_ *< π*) or (*θ*_*stim*1_, *θ*_*stim*2_ *> π*). ReactCategoryAnti: Match if opposite category (*θ*_*stim*1_ *< π* & *θ*_*stim*2_ *> π*) or (*θstim*1 *> π* & *θstim*2 *< π*).

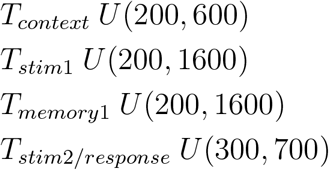

### 1.3 Training procedure

The loss *L* to be minimized was computed by time-averaging the squared errors between the network output **z**(**t**) and the target output 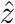(*t*).

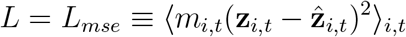

Here, *i* was the index of the output units and *t* was the index for time. We implemented a mask, *m*_*i,t*_, for modulating the loss with respect to certain time intervals. For example, in the first 100 ms of the context and response periods there was a grace period with *m*_*i,t*_ = 0. During the response period, *m*_*i,t*_ = 5 and for the rest of the trial *m*_*i,t*_ = 1. For the fixation output unit, *m*_*i,t*_ was two times stronger than the mask for the *ϕ*_*response*_ output units. The training was performed with Adam, a variant of stochastic gradient descent [2]. The learning rate ranged from 10^*-*4^ (tanh networks) to 10^*-*3^ (all other networks). The decay rate for the first and second moment estimates were 0.9 and 0.999, respectively. During training, we randomly interleaved all the tasks with equal probabilities, except for the ContextIntModality1 and ContextIntModality2 tasks that appeared five times more frequently to prevent the network from integrating both modalities equally. This alternative strategy gave the network an accuracy close to 75% if trials from these tasks were not over-represented. During training, we used mini-batches of 64 trials, in which all trials were generated from the same task for computational efficiency. Training was terminated when *L* stopped decreasing, which was generally after 5e7 training steps.

The network and training were implemented in TensorFlow.

### 1.4 Alternative hyperparameters and network architectures

We trained networks with the following possible hyperparameters and architectures:

The network architecture was either the leaky RNN architecture defined previously, or the leaky GRU architecture [3].

We explored a number of nonlinear functions *σ*(·),

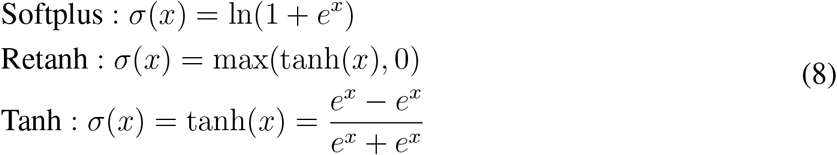

We initialized each weight matrix from a diagonal matrix,

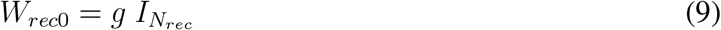

or from a random Gaussian

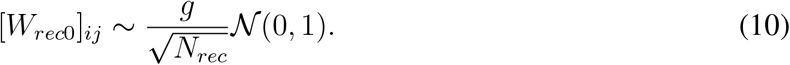

Here, *g* scaled the values in the initial weights. In networks with the tanh activation function and the leaky RNN architecture *g* = 1 and all other networks *g* = 0.8. We found that networks with the tanh activation function required this higher *g* value to prevent quenching network activity during training.

To avoid overly complex solutions that didn’t generalize well, we penalized high activity and strong weights using an L2 regularization on **h** and on each weight matrix *W*_*in*_, *W*_*rec*_, *W*_*out*_. The hyperpa-rameter selection criterion was the highest level of regularization that resulted in >80% performance on held out test data. We found the highest level regularization that still resulted in greater than 80% performance on all tasks to be 10^*-*6^ for both weight and activity regularization.

### 1.5 Fixed Points

Our networks were high-dimensional nonlinear systems, rendering them difficult to understand intuitively. Examination of these networks was made easier through analysis of fixed points, which are locations in state space where the motion of the system is approximately zero. Through a Taylor expansion of our dynamical equation, we may see that our nonlinear system can be approximated as a linear one around fixed points, **h**^*∗*^:

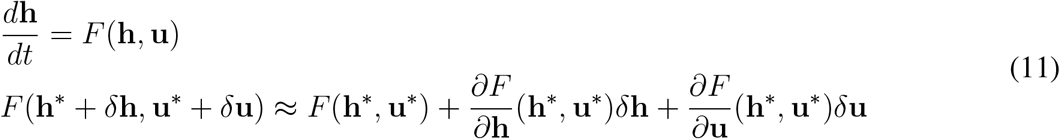

The second order terms (not shown) are approximately zero because 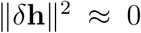. The first term, F(**h**^*∗*^,**u**^*∗*^) is zero by definition of the fixed point, where **h**^*∗*^ is the location where the update *F* (**h**^*∗*^, **u**^*∗*^) = 0. For the majority of this work, with the exception of Supplementary Figure 6, we hold input values to their constant value during a task period. We can therefore ignore the last term where *d***u** = 0. Therefore, around fixed points, we can approximate our nonlinear dynamical systems as the linear system, 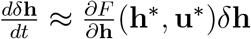.

Eigendecomposition of the matrix, 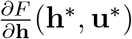, reveals in which dimensions of state space the dynamics are contracting, expanding or are marginally stable (ie. neither contracting or expanding).

Eigenvectors with an associated real part of the eigenvalue *λ <* 1 are contracting dimensions, *λ >* 1 are associated with expanding dimensions and *λ* ≈ 1 are marginally stable. At a fixed point that is contracting in every dimension, the state is at a basin of attraction. This is a particularly useful dynamical system for preparing an optimal initial condition for the next task period. Marginally stable dimensions are useful for integrating noisy pulses of stimulus information and for memory of a continuous variable. Saddle points are contracting in some dimensions, while repulsive in other dimensions. Saddle points are useful for decision making along the repulsive dimensions. Repulsive dimensions can additionally be useful for keeping the neural state away from a particular region of state space. For example, the neural state must remain outside of output potent space (orthogonal to the readout weights) until the response period.

To identify fixed points, we empirically optimized for a set of {**h**^**1***∗*^, **h**^**2***∗*^, …} satisfying *F* (**h**^*∗*^, **u**^*∗*^) = 0 while defining **u**^*∗*^ by holding the inputs **u** constant for each task period. Each different input condition reconfigured the RNN into a new dynamical landscape with a different set of fixed points. Therefore, each set of {**h**^**1***∗*^, **h**^**2***∗*^, …} was associated with a particular input **u**^*∗*^. At the fixed point **h**^*∗*^ with inputs **u**^*∗*^, the update to the state at the next time point is zero and therefore, the state doesn’t move away from this location. We used the term fixed point to include approximate fixed points, where the update is small on the timescale of our task. Our fixed points range between *q* = 10^*-*3^ to 10^*-*15^ where 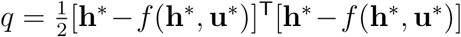 We included a wide range of *q* values in order to best highlight relevant dynamics on a case by case basis.

We used the Fixed Point Finder package in Tensorflow [4].

### 1.6 Input Interpolation

We examined how fixed point structures moved and changed stability as the dynamical system was reconfigured by different inputs. To do this, we interpolated across pairs of inputs and identified fixed points for each intermediate input condition. For input vectors **u**_**1**_ and **u**_**2**_, we identified fixed points for *α***u**_**1**_ + (1 - *α*)**u**_**2**_ where *α* was varied between 0 and 1 in 0.05 increments.

### 1.7 Analysis of Fixed Points for Interpolated Inputs

After input interpolation [see Section 1.6 for details], we wanted to compare fixed points across input conditions in order to track their positions and stability in high dimensional space. However, there were often multiple fixed points, making it difficult to track an individual fixed point across input conditions. We focused on the fixed point closest to the state at the end of a task period of interest (except in Figure 4f, where we focused on the closest unstable fixed point because it appeared more relevant to the nonlinear dynamics - the closest fixed point was stable and was also shared across tasks). Our reasoning was that if the state evolved toward a particular fixed point, it was likely relevant for computation. After identifying fixed points during rule input interpolation, we ran the network forward from the beginning of the context period for each interpolated rule input and identified the fixed point closest to the network state at the end of the task period of interest (stimulus or response period). We refer to this closest fixed point as the ‘relevant’ fixed point for a given interpolated input. We calculated the Euclidean distance between relevant fixed points associated with adjacent interpolated input conditions (*α*_1_, *α*_2_) as:

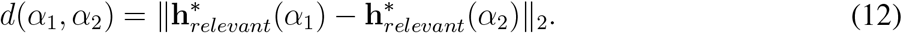

We also tracked the stability of the relevant fixed point for each interpolated input. To do this, we performed eigenvalue decomposition on the Jacobian of the RNN state transition function at the relevant fixed point,

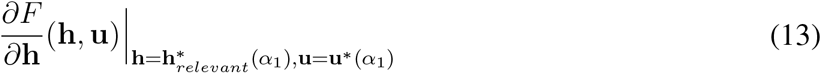

The eigenvalue with the maximum real value is informative about whether the relevant fixed point is stable. By tracking the stability over input interpolation, we could identify bifurcations in the dynamical landscape.

To examine the relevant dynamical motif for a given task period, we defined a ‘relevant fixed point’ to be the fixed point closest to the state at the end of the task period. If the input interpolation between *α* = 0 and *α* = 1 resulted in approximately the same location of the relevant fixed point and approximately the same local dynamics around the relevant fixed point, then we defined the relevant fixed point as being functionally the same across inputs; and therefore, the dynamical motif was shared across input conditions.

Alternatively, if the interpolation between *α* = 0 and *α* = 1 resulted in a bifurcation of the fixed point structure, then we defined the dynamical motifs to be distinct. We highlight that our definition of distinct motifs was limited in that a different path for consecutive input interpolation might not result in a bifurcation. It will be of great interest to explore ambiguous cases of shared and distinct motifs in future work.

### 1.8 Effective Input Modulation

In previous work, it was identified that relaxation dynamics of the network state could contextually integrate stimulus inputs [5]. We identified some networks that additionally contextually amplified stimulus inputs (Supplementary Figure 6). In order to deconstruct how the signal from one input is contextually amplified, we look at the first order Taylor Series approximation of the state update around a particular input, **u**^*∗*^, and its associated fixed point, **h**^*∗*^:

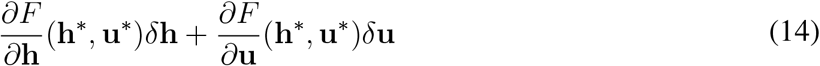

The network received contextual rule input during the context period and moved toward a stable fixed point. We took this stable fixed point during the context period to be the initial conditions for the subsequent stimulus period. We modelled the initial stimulus input in the ▵*u*_*stimulus*_ = *u*_*stimulus*_ *u*_*context*_ term without changing 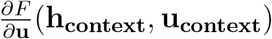 at the context dependent fixed point. We calculate the input response for each stimulus condition for trials spanning [0, 2*π*) by calculating the norm of the dot product of

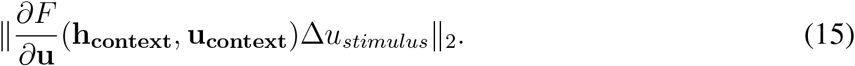

We found that there was task specific amplification of stimulus modality inputs, where the modality 1 (2) input response was larger for tasks where the network must respond according to modality 1 (2).

### 1.9 Task variance analysis

In order to examine the contributions of unit variance to computation in each task period, we used a modified version of the task variance analysis described previously [1]. We ran the network forward for a set of possible stimulus conditions on the task of interest. For example, in the Delayed Response tasks, we presented the network with trials where *θ*_*stimulus*_ ranged from [0, 2*π*). In the Decision Making tasks we ran the same network with with *θ*_*stimulus*_ ranging from [0, 2*π*) and coherences ranging from 0.005 to 0.2. We then computed the variance across possible stimulus conditions for each unit on each task period through time. This was a deviation from [1] in two ways. Firstly, we computed task period variance across stimulus conditions for all task periods separately because we considered each task period as a separate dynamical landscape. Secondly, we computed variance through time rather than averaging across time, because we were interested in the dynamics rather than static representations. Variance during the fixation period was low so we excluded the fixation period from this analysis, and we study it separately in Figure 4b-c. Private noise (*ξ* in equation 4) was set to zero for this analysis to eliminate the effect of recurrent noise. This analysis was a useful method to uncover unit contributions to network computations because our networks were activity regularized. Given activity regularization, any deviations from zero activity were costly for the network to produce and therefore likely beneficial for task computation. The result of this analysis was a matrix composed of columns of units and rows of task periods, where each index quantified the participation of a given unit to the computation during a given task period. We refer to this matrix as the variance matrix.

Correlations between variance matrices were computed by first sorting task period rows according to one reference network. A correlation matrix for each network was computed by finding the Pearson correlation between rows of the variance matrix for that network separately. Each correlation matrix was compared to every other correlation matrix by first flattening the the upper triangle of entries in each correlation matrix. We then calculated the Pearson Correlation between this vector and the same vector associated with each trained network. Both trained and untrained networks were compared to trained networks in order to determine whether the structure in trained networks emerged due to the input structure or due to learned dynamical motifs.

### 1.10 Clusters

We sorted rows and columns of the variance matrix [see Section 1.9 for details] according to similarity using the Ward variance minimization algorithm [6]. This algorithm produced a dendrogram that shows the hierarchical distance between rows or columns of the task variance matrix. In order to obtain discrete clusters, we identified the optimal distance threshold for each dendrogram by computing the silhouette score on the basis of intracluster and intercluster distances. The silhouette score of an unit i was 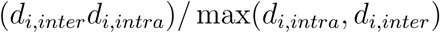, where *d*_*i,intra*_ was the average distance of this unit with other units in the same cluster, and *d*_*i,inter*_ was average distance between this unit and units in the nearest cluster. The silhouette score of a clustering scheme was the average silhouette score of all units. A higher silhouette score meant a better clustering. We computed the silhouette score for the number of clusters ranging from 3 to 40. The optimal number of clusters k was determined by choosing the k with the highest silhouette score. Clustering results were robust to clustering method and to the network hyperparameters that we explored.

### 1.11 Lesions

We lesioned a network unit by setting its projection weights to 0 for all recurrent and output units. When we lesioned a particular network cluster, we lesioned all units within that cluster.

### 1.12 Transfer Learning

Networks were pre-trained on a subset of tasks as described previously, where *W*_*in*_, *W*_*rec*_, *W*_*out*_ and bias vectors **b**_**in**_, **b**_**out**_ were learned over the course of training [see Section 1.3 for details]. After this initial stage of training, the network was trained on a held out task. In this second phase of training, all network connections were held fixed except for the rule input weights of the held out task. That meant that in the second phase of learning, only a vector **u**_**rule**_*∗* of size *N*_*rec*_ within *W*_*in*_ changed.

In Supplementary Figure 7, we wanted to understand the relationship between how well this transfer learning approach worked and whether the held out task required learning of novel dynamical motifs. We first quantified the extent to which a task required a unique dynamical motif by comparing rows in the variance matrix [see Section 1.9 for details]. We sorted rows according to a reference network and computed the correlation matrix for the variance atlas in each network across all hyperparameter settings in Figure 3b. We then took the average across all correlation matrices. We used this average correlation matrix to inform the average relationship across task periods for all networks that we examined. For each task, we identified the maximum correlation to other tasks for each task period. The most unique task period was that which had the lowest maximum correlation to other task periods. Our hypothesis was that tasks with lower correlation required unique dynamical motifs whereas tasks with higher correlation could be shared across tasks. We then quantified how well our transfer learning method performed for each task. We first trained a network on all but one task in the first stage of learning and then on the held out task in the second stage of learning. We compared the cost during training each task in the second stage of transfer learning to a single task network. Single task networks were trained to perform the task of interest with all weights and biases plastic. We compared the cost at two different points in the training process, (1) early in training in order to determine the benefit from starting with previously learned dynamical motifs and (2) late in training to determine the cost of freezing all weights except the rule input for the task of interest. We then plotted the difference in cost at these two separate time points against our metric for unique task periods. The fast learning benefit was smaller and the long term cost was negative for tasks with unique dynamical motifs.

## Supplementary Figures

**Supplementary Figure 1:**
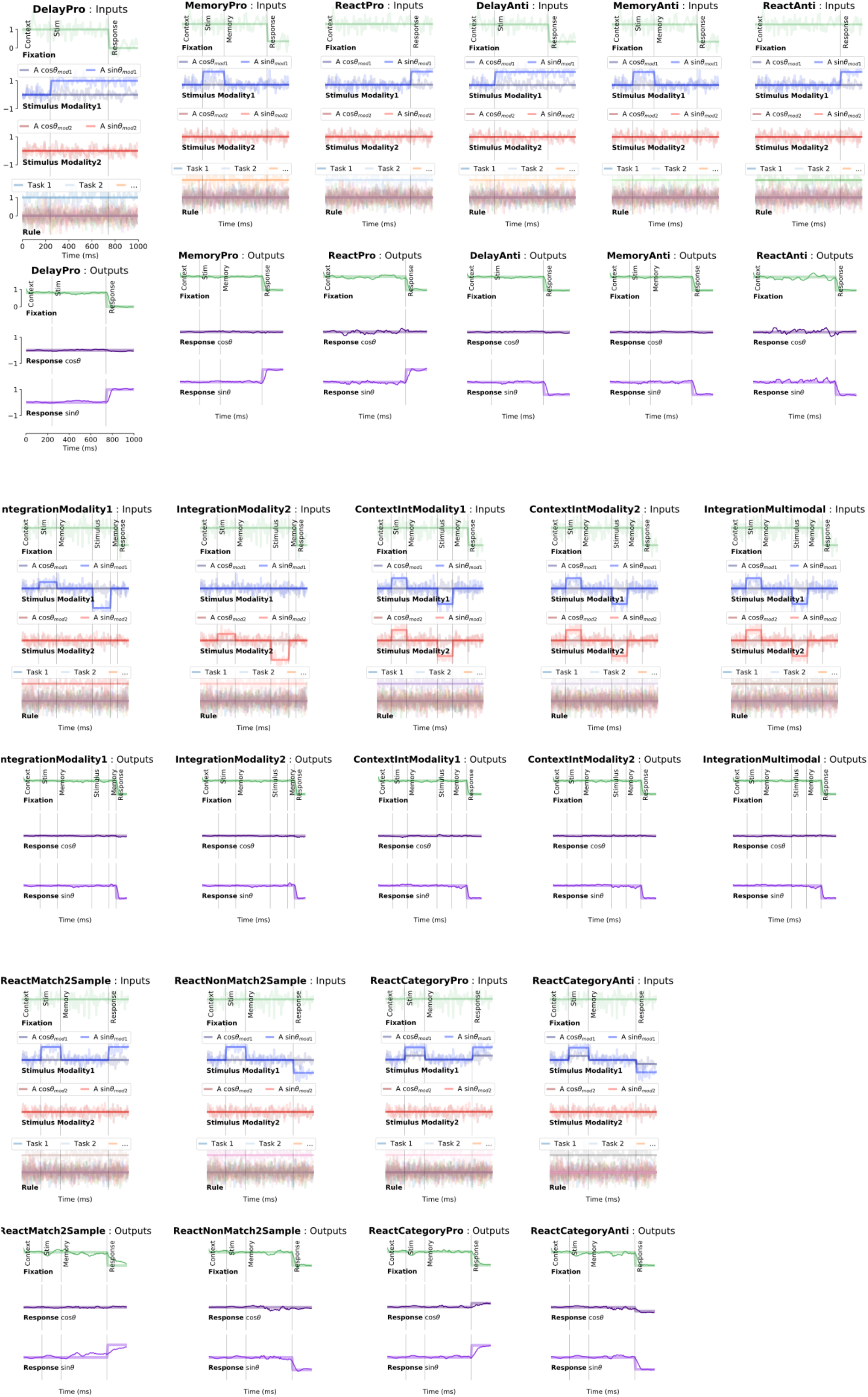
Inputs and outputs for each task for one example trial. One panel for each task. **Top:** Noisy fixation, stimulus (modality 1 and 2), and rule input time-series (overlayed without noise for clarity.) Noise was used during training while analyses were performed on running the network without noise. Vertical lines divide task periods. **Bottom:** Targets (thick lines) overlaid with outputs of a trained network (thin lines).

**Supplementary Figure 2:**
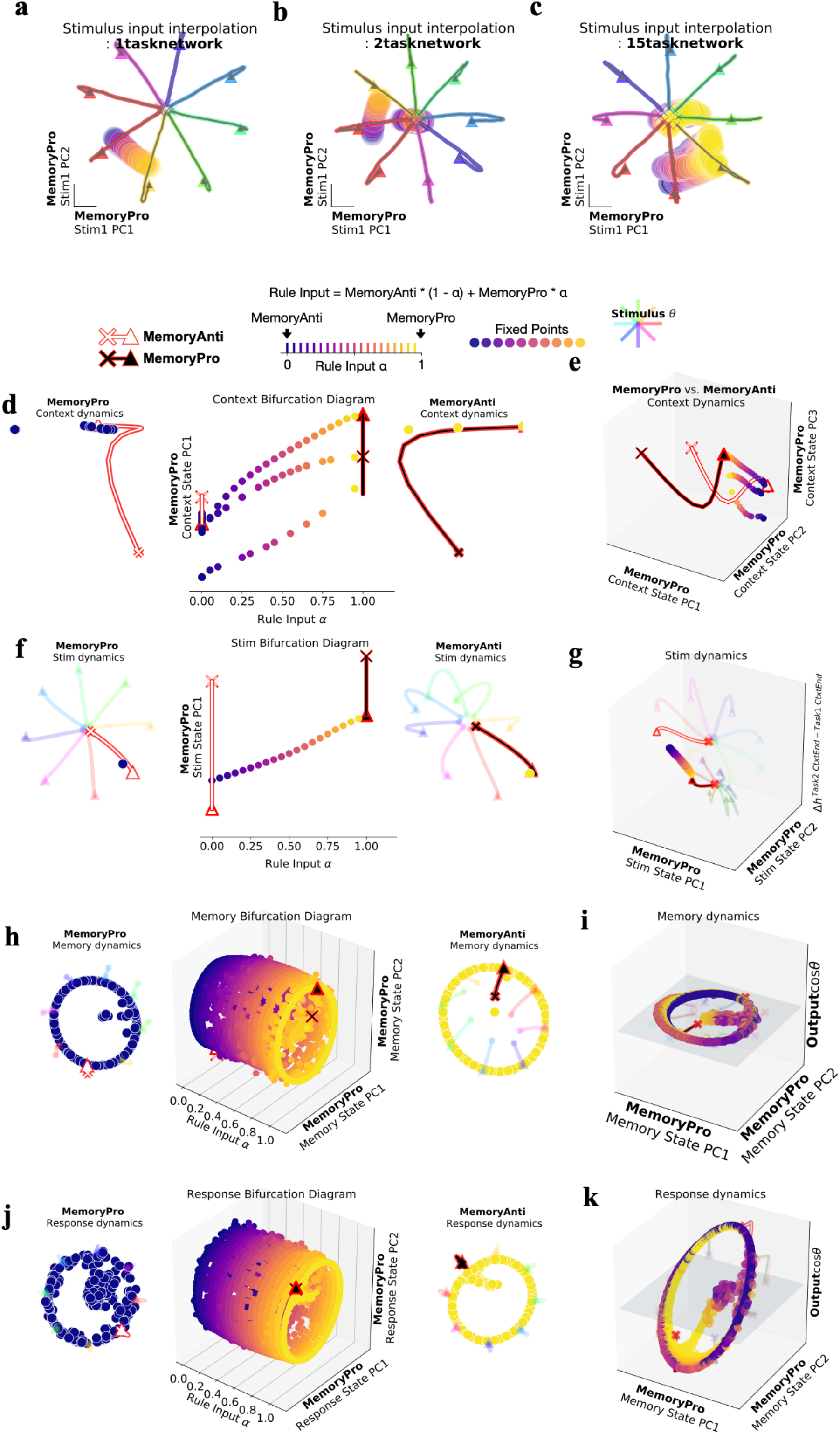
Fixed point structure is dependent on other tasks the network is trained to perform. **(a)** Fixed points for interpolated inputs across stimulus conditions 0°(blue)→45° (yellow), reveals one fixed point that moves dependent on stimulus input conditions. Trajectories colored according to stimulus orientation. Additional fixed points are revealed in **(b)** 2 task networks and **(c)** 15 task networks. **(d-k)** *N*_*rec*_ *= 1024* network with softplus activation, diagonal initialization and no input noise or private noise (see equations 2 and 7 in Methods for noise definitions). Fixed points for interpolation between inputs for MemoryAnti (*α = 0*) and MemoryPro (*α = 1*) tasks during **(d**,**e)** context **(f**,**g)** stimulus **(h**,**i)** memory **(j**,**k)** and response periods. **(d) middle:** Fixed points for 20 intermediate *α* values (x-axis) projected into top PC defined by state evolution during the context period of the MemoryPro task (y-axis) with (**left)** MemoryPro *α = 0* and **(right)** MemoryAnti *α = 1* fixed points and trajectories. **(e)** Fixed points for rule input-interpolation between tasks, MemoryPro (blue fixed points, white state trajectory) and MemoryAnti (yellow fixed points, black state trajectory) projected into the top three MemoryPro context period state evolution PCs. **(f)** Same as **d** for stimulus period, with fixed points projected into top PC defined by the state evolution during the stimulus period of the MemoryPro task (y-axis) **(g)** Same as **e** for stimulus period, projected into the top two MemoryPro stimulus period state evolution PCs (x and y axes) and the dimension separating both tasks at the end of the context period (z axis). **(h)** Same as **d** for memory period, projected into top two PCs defined by the state evolution during the memory period of the MemoryPro task (y and z-axes) **(i)** Same as **e** for memory period, projected into the top two MemoryPro memory period state evolution PCs (x and y-axes) and the output weight vector (from *W*_*out*_) associated with *cosθ*_*stimulus*_ on the z-axis. **(j)** Same as **d** for response period, projected into top two PCs defined by the state evolution during the response period of the MemoryPro task (y and z-axes) **(k)** Same as i for response period.

**Supplementary Figure 3:**
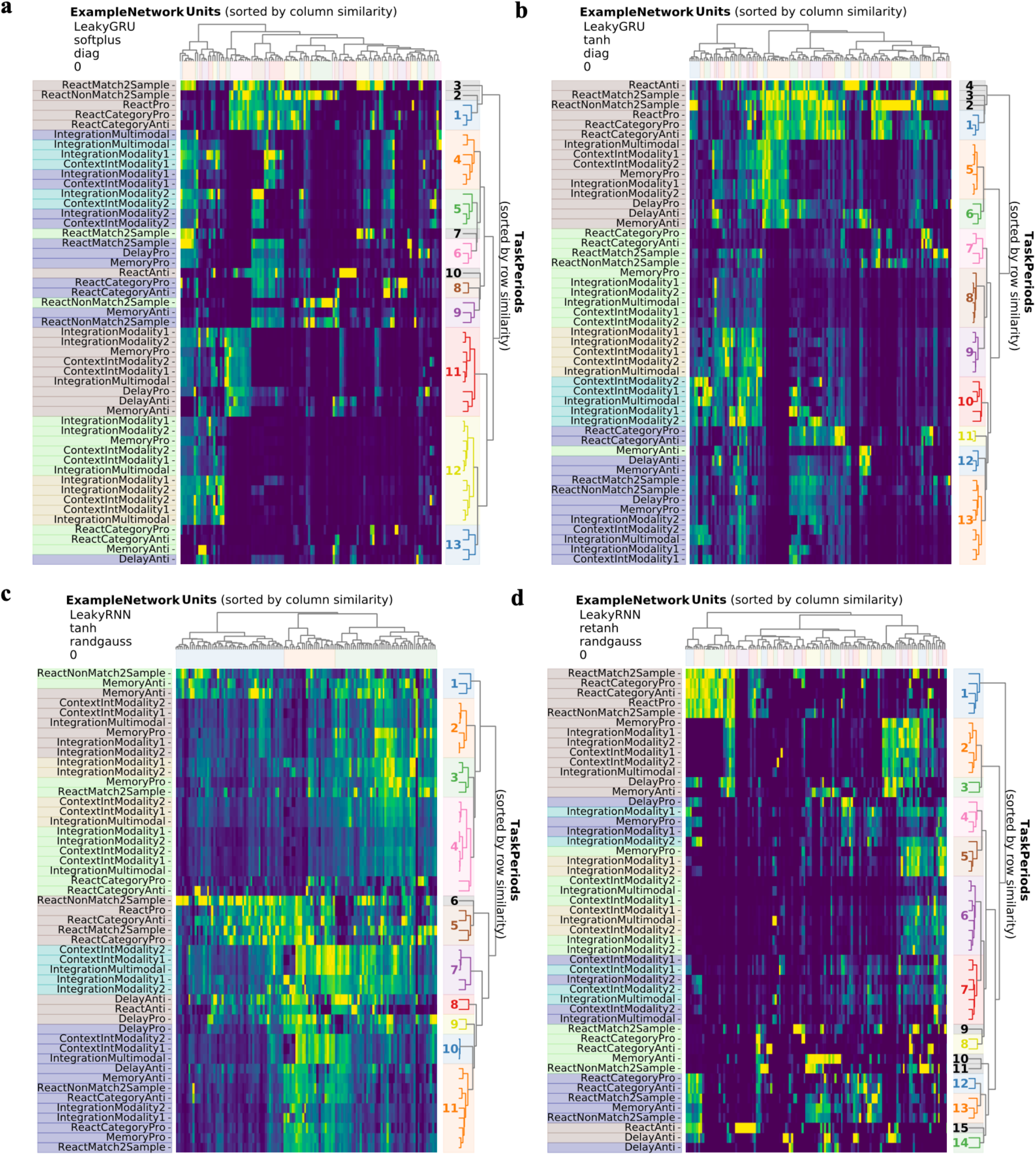
Example variance matrices for four different networks. Variance matrix: variance of unit activations across stimulus conditions normalized across task periods. Rows and columns sorted according to similarity (see Methods Section 1.10). **(a)** LeakyGRU, softplus activation, diagonal initialization, *N*_*rec*_*=128* **(b)** LeakyGRU, tanh activation, diagonal initialization, *N*_*rec*_*=128* **(c)** LeakyRNN, tanh activation, random Gaussian initialization, *N*_*rec*_*=128* **(d)** LeakyRNN, retanh activation, random Gaussian initialization, *N*_*rec*_*=128*.

**Supplementary Figure 4:**
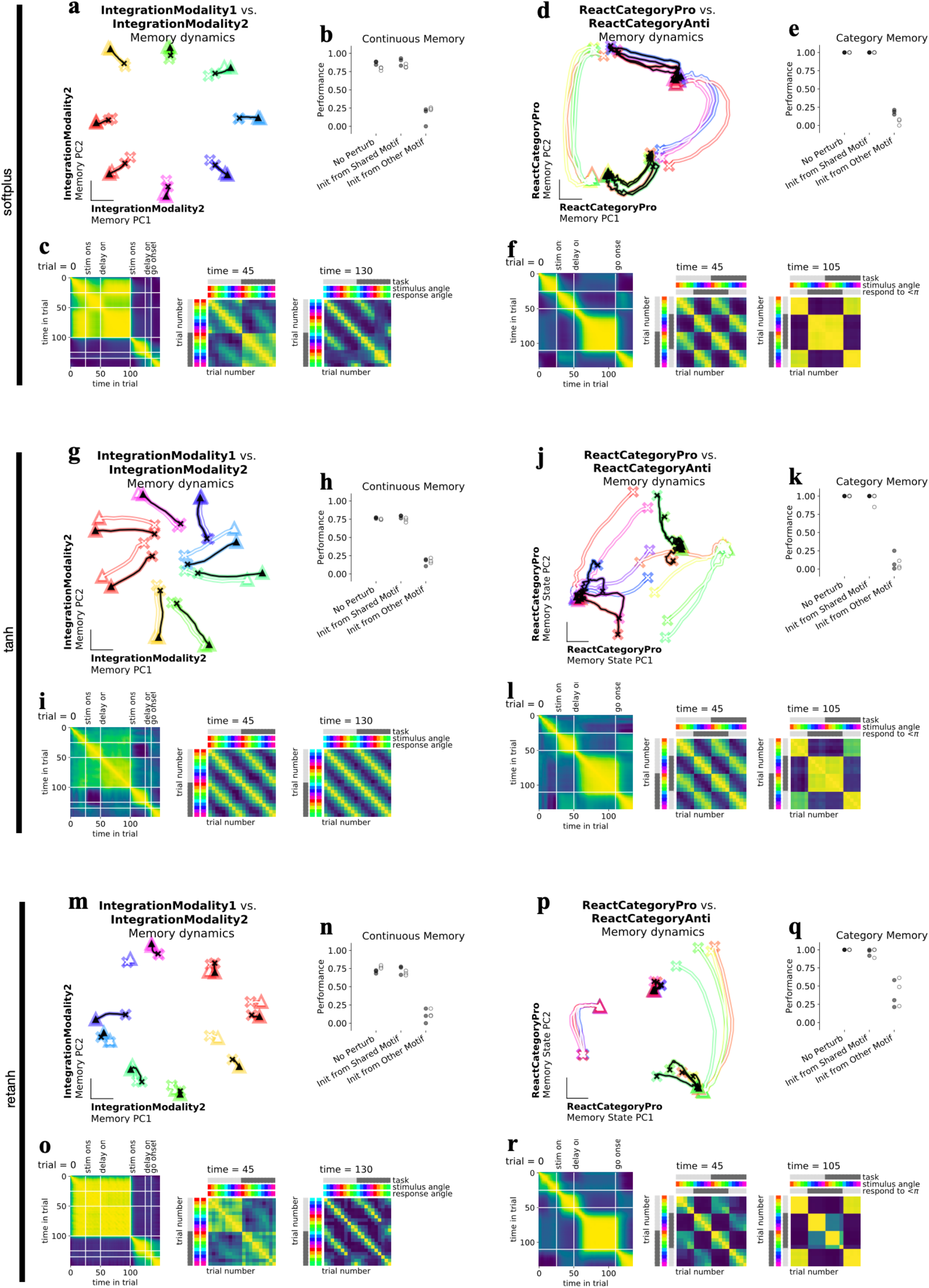
Discrete and continuous variable memory computations result in two kinds of shared memory attractors. Networks were trained with softplus activation function for **a-f (a)** Neural state trajectories for 8 stimulus conditions (colored by response direction) starting from ‘x’ in IntegrationModality2 PC space for IntegrationModality1 (white) and IntegrationModality2 (black) tasks. **(b)** Perturb initial condition for IntegrationModality1 response period state to the memory period final state of either IntegrationModality2 task (Init from shared motif), ReactCategoryAnti (Init from other motif) or IntegrationModality1 (No perturb) for trials with the same response direction and run the network forward through the response period. One data point for each of three networks trained with diagonal initialization (filled circles) or with a random gaussian initialization (open circles). Performance was similar across ‘No perturb’ and ‘Init from shared motif’ conditions but was dramatically impacted for ‘Init from other motif’ conditions. **(c) left:** Correlation across timesteps throughout an IntegrationModality1 trial. Task period transitions highlighted in white lines. **middle:** Correlation across trials for different stimulus conditions for IntegrationModality1 and IntegrationModality2 tasks at the end of the first stimulus period. **right:** Correlation across trials for different stimulus conditions for IntegrationModality1 and IntegrationModality2 tasks at the end of the memory period. Trials are highly correlated across tasks according to response direction. **(d)** Same as (a) for two category memory tasks, ReactCategoryPro (white) and ReactCategoryAnti (black). **(e)** Perturb initial condition for ReactCategoryPro response period state to the memory period final state of ReactCategoryAnti task (Init from shared motif), IntegrationModality1 (Init from other motif) or ReactCategoryPro (No perturb) for trials with the same response direction and run the network forward through the response period. **(f) left:** Correlation across timesteps throughout an ReactCategoryPro trial. Task period transitions highlighted in white lines. **middle:** Correlation across trials for different stimulus conditions for ReactCategoryPro and ReactCategoryAnti tasks at the end of the stimulus period. **right:** Correlation across trials for different stimulus conditions for ReactCategoryPro and ReactCategoryAnti tasks at the end of the memory period. Trials are highly correlated across tasks according to response direction. **(g-l)** Same as (a-f) for tanh activation function. **(m-r)** Same as (a-f) for retanh activation function.

**Supplementary Figure 5:**
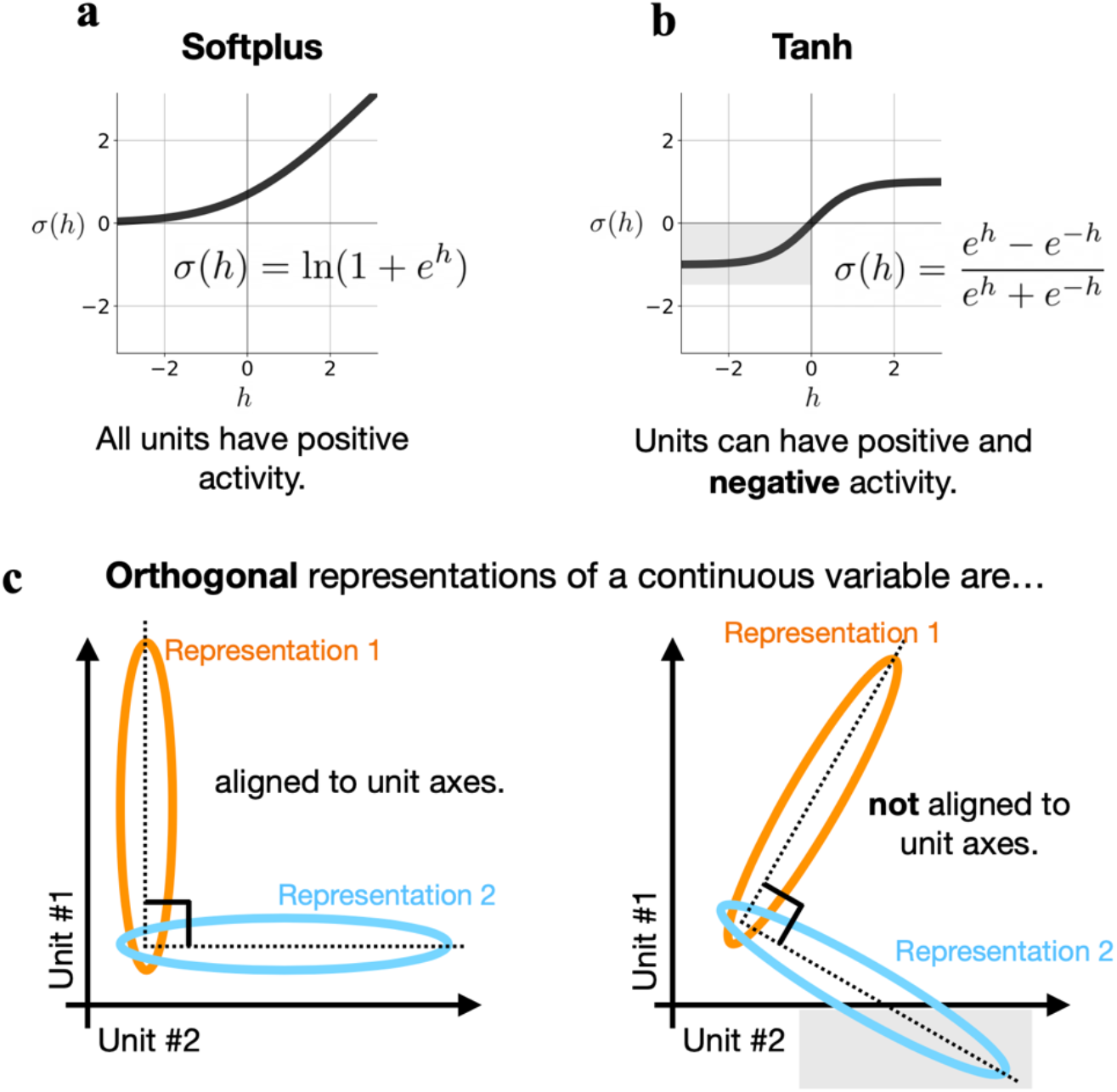
Positive activation functions align orthogonal motifs to unit axes. **(a)** Softplus activation function maps activity to the positive real line. **(b)** Tanh activation function does not impose positivity constraints. **(c) left**: Two activity patterns that are constrained to lie in the positive orthant can only be orthogonal to one another when aligned to the unit axes. **right:** Activation functions without positivity constraints do not constrain activity patterns to unit axes. *Note: Either system can use additional dimensions to orthogonalize representations not aligned to unit axes in an infinitely large network*.

**Supplementary Figure 6:**
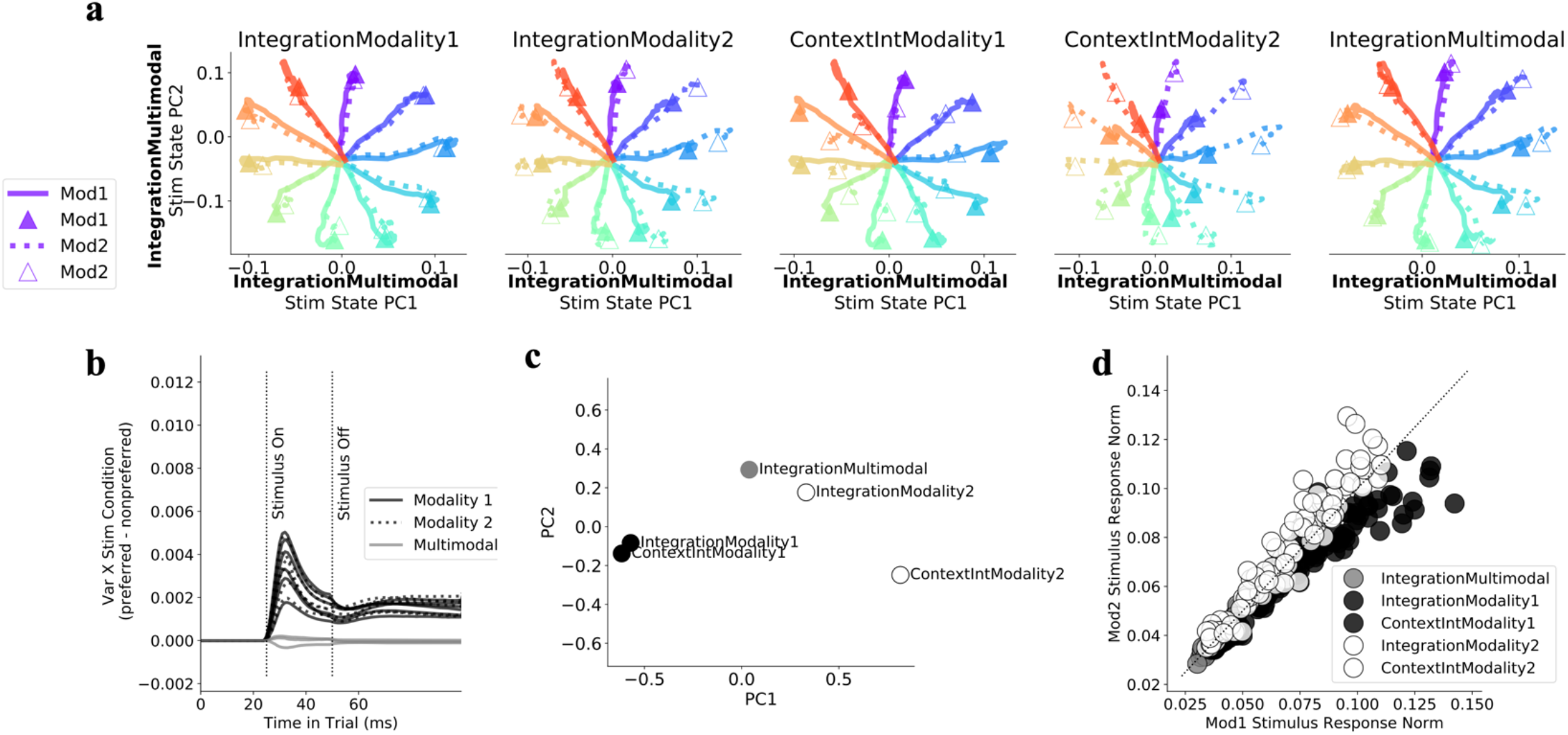
Input modulation for context dependent integration. **(a)** Stimulus dependent network state for both modality 1 (solid) and 2 (dotted) for 5 different tasks where relevant modality is specified in title for each subpanel. **(b)** Difference between variance across stimulus conditions for preferred modality minus variance across stimulus conditions for nonpreferred modality. Preference is defined by task rules. One line for each of five tasks in three different networks. **(c)** Context period network state for all 5 tasks projected onto the first and second PCs defined by variance of the state for each task. (PCA on 5xN_rec_ matrix). Organization of network states at the end of the context period shows compositional structure. PC1 goes from modality 1->2 and PC2 separates ContextIntegration vs. Integration decision tasks. **(d)** Norm of input jacobian for 24 stimulus conditions spanning [0, 2fi) on each of 5 tasks (see Methods Section 1.8). Trials where the network should prefer modality 1 are black, modality 2 white, multimodal gray. Hyperparameters: LeakyRNN, softplus activation, diagonal initialization, *N*_*rec*_*=128*

**Supplementary Figure 7:**
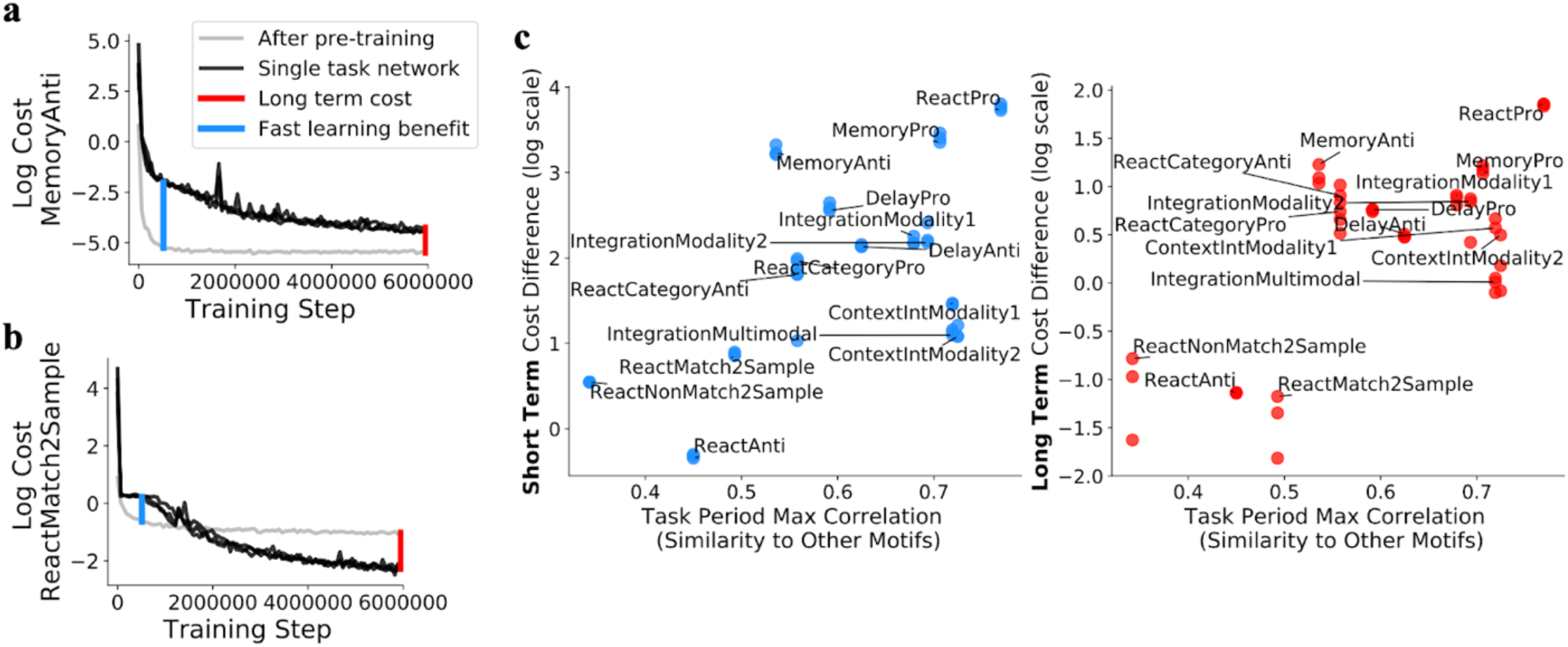
Leave-one-out transfer learning is less effective for tasks with unique dynamical motifs. **(a)** Log cost during training single rule input weights for **(a)** MemoryAnti and **(b)** ReactMatch2Sample tasks after pre-training all other weights on all other tasks (gray) and during training all weights and biases on a single rule (black, 3 lines for 3 different networks with different random reeds). Single task networks provide a baseline comparison to measure how well the pre-training method works for any given task. **(c)** Cost difference between single task networks and transfer learning network (as in a,b) plotted against task period max correlation (See Methods Section 1.12 for details) early in learning (left) and late in learning (right) for each task. The fast learning benefit was smaller and the long term cost was negative (single task networks outperformed transfer learning networks) for tasks with unique dynamical motifs. Hyperparameters: LeakyRNN, softplus activation, diagonal initialization, *N*_*rec*_*=128* for transfer learning and single task networks. Task period max correlation was averaged across networks with all hyperparameter settings in Fig.3b.

